# PSC niche develops into immune-responsive blood cells capable of transdifferentiating into lamellocytes in *Drosophila*

**DOI:** 10.1101/2022.10.17.512558

**Authors:** Alexander Hirschhäuser, Darius Molitor, Gabriela Salinas, Jörg Großhans, Katja Rust, Sven Bogdan

## Abstract

*Drosophila* blood cells called hemocytes form an efficient barrier against infections and tissue damage. During metamorphosis, hemocytes undergo tremendous changes in their shape and behavior preparing them for tissue clearance. Yet, the diversity and functional plasticity of pupal blood cells have not been explored. Here, we combine single-cell transcriptomics and high-resolution microscopy to dissect the heterogeneity and plasticity of pupal hemocytes. We identified precursor and effector hemocytes with distinct molecular signatures and cellular functions clearly distinct from other stages of hematopoiesis. Strikingly, we identified that PSC cells, which function as lymph gland niche, are highly migratory and immune responsive cells in the pupa. PSC cells can transdifferentiate to lamellocytes triggered by wasp infection. Altogether, our data highlight a remarkable cell heterogeneity, and identifies a cell population that acts not only as a stem cell niche in larval hematopoiesis, but functions as cell reservoir to pupal and adult blood cells.

## Introduction

The innate immune system depends on a wide array of conserved cellular and molecular strategies to mediated pathogen defense, tissue remodeling and repair. *Drosophila* is a powerful genetic model organism to study blood cell development and innate immunity ^1, 2^. The fruit fly has an open circulatory system in which the heart pumps blood, the so-called hemolymph into the body cavity circulating all organs. To fight against infections, flies evolved a large variety of defense responses that share highly conserved features with the human innate immunity ^3, 4^. Similar to mammals, the first line of defense against invading pathogens and wounds in *Drosophila* relies on both, a humeral response by which effector molecules such as antimicrobial peptides are secreted into the hemolymph and a cellular response, in which pathogens are phagocytosed by blood cells, the so-called hemocytes ^5, 6^. Hemocytes have been traditionally classified by their cell morphology into three different effector cells, plasmatocytes, crystal cells, and lamellocytes ^7^. Plasmatocytes, the most abundant immune cell type in flies, are professional phagocytes similar to mammalian bone marrow-derived macrophages ^6^. Circulating and tissue-resident plasmatocytes are immediately recruited to sites of wounding and infections and mediate the major cellular immune response by phagocyting pathogens and secreting antimicrobial and clotting factors ^8, 9^. Upon injury, platelet-like crystal cells are required for the melanization of wounds, a process that involves the rapid synthesis of the black pigment melanin required for wound healing and encapsulation of invading parasites. Lamellocytes, by contrast, are rarely observed in healthy flies, but are dramatically induced in response to infection by parasitic wasps ^10^.

Blood-lineage specification in flies requires a similar conserved set of transcriptional regulators and signaling pathways to those that control mammalian hematopoiesis including the GATA factor Serpent (Srp) and the friend of GATA (FOG) transcription factor U-shaped which together determine blood cell fate ^11^. *Drosophila* hematopoiesis occurs in two spatially and temporally distinct phases with clear parallels to mammals^12^. Blood cells initially derive from head mesoderm of the developing embryo and give rise to both plasmatocytes and crystal cells. These cells colonize and self-renew in segmentally repeated epidermal-muscular niches in larvae ^13^. The hematopoietic pockets provide both an attractive and trophic microenvironment, promoting proliferation of the initial 600-700 embryonic cells to about 9000-10000 cells per third-instar in the differentiated state ^14^. The second wave of hematopoiesis in flies occurs post-embryonically in the lymph gland of larvae, a specialized hematopoietic organ of mesoderm origin, which is arranged in multiple paired lobes along the anterior part of the dorsal aorta ^15^. The lymph gland harbors progenitors, differentiating and mature blood cells within distinct zones, the cortical zone with mature hemocytes, the medullary zone (MZ) with progenitors and the posterior signaling center (PSC). PSCs are thought to function as a stem cell niche to control the differentiation of all three effector types from progenitors ^16^. The PSC comprises a small cluster of about 30-40 cells and are mitotically inactive in the mature lymph gland. These cells are marked by the expression of the homeobox protein Antennapedia (Antp) and high levels of the conserved member of the Early B-cell Factor (EBF) family of transcription factors, Collier/Knot (Col/Kn) that controls PSC specification and cell numbers ^17, 18^. However, genetic ablation experiments showed that the PSC is dispensable for blood cell progenitor maintenance and further revealed a key role of the conserved EBF transcription factor Col/Kn as an intrinsic regulator of hematopoietic progenitor fate ^19^. To date, the importance and function of PSC cells after pupal lymph gland histolysis is unclear but the recent FlyCellAtlas identified PSCs in adult flies, suggesting that PSCs are important beyond lymph gland niche function ^20^.

Under normal conditions, progenitors in the lymph gland give rise to plasmatocytes and crystal cells. Both effector cell types are released into the circulation as the lymph gland dissociates at the onset of metamorphosis ^21^. Thus, with the onset of pupation, both embryo and lymph gland-derived blood cells mix together and populate the pupa. Recent advances in single-cell RNA sequencing (scRNA) technologies further revealed a much larger diversity of the hemocytes present in the *Drosophila* larva as well as in the lymph gland ^22–24^. Comparative analysis of these datasets showed important differences in the molecular signatures between hemocytes during embryogenesis and larval stages and further suggest an even increased complexity of hemocyte populations in metamorphosis and in adult flies ^20, 25, 26^. Metamorphosis goes along with dramatic and systemic physiological changes that requires an integration of the innate immune system. In response to ecdysone, hemocytes rapidly upregulate cell motility and phagocytosis of apoptotic debris, and acquire the ability to chemotax to tissue damage. ^9, 27, 28^. Bulk RNA-seq gene expression analysis recently uncovered thousands of genes that are differentially expressed in pupal hemocytes compared to larvae ^29^. However, whether this striking differential expression pattern of pupal hemocytes might also reflect the differentiation of new hemocyte subtypes or cell types has not yet been addressed. Insights into heterogeneity of these activated hemocytes would, however, also shed light on how tissue clearance is regulated during development.

Here, we employ the scRNA seq technology to characterize the molecular signatures of pupal hemocyte subpopulations. Our single-cell data reveals the presence of different precursor and effector hemocytes with distinct transcriptomic signatures and cellular functions including proliferative potential, antimicrobial peptide production, proteasomal degradation, fatty acid oxidation, and phagocytosis. Cross-stage dataset analysis provides evidence that most pupal hemocyte populations display distinct transcriptomic profiles suggesting specific functions during metamorphosis. Among the pupal blood cell types, we identified a highly migratory and immune responsive PSC cell cluster that persist into the adult fly. Lineage tracing experiments analysis reveal that these plastic PSC cells are able to differentiate to lamellocytes in response to wasp infection. This displays the remarkable ability of a niche cell to transdifferentiate in response to environmental cues.

## Results

### *Drosophila* pupal hemocytes display remarkable cellular heterogeneity

Recent comparative scRNA-seq analyses revealed cellular heterogeneity in the molecular signatures of *Drosophila* hemocytes but also important differences during embryogenesis and larval stages ^22–24, 26^. Both, embryo and lymph gland-released hemocytes mix together at the onset of pupariation and persist into the adulthood. Our recent bulk RNA analysis already revealed highly differential expression profile in pupal hemocytes compared to larvae ^29^. To analyze this mixed blood cell population in more detail, we used scRNA-seq technology to dissect the cellular heterogeneity of the total blood cells in *Drosophila* early pupa. We applied a high-throughput full coverage of transcripts approach using the SMART-Seq technology on the ICELL8 system (Figure 1A). ^30^. The use of the ICELL8 platform allows beside the use of flexible chemistry for library preparations on a one-nanowell chip reaction the analysis of living or fixed cells with a large range of sizes ^30^. Hemocytes were isolated from 2-10h APF pupae. Wild type live or fixed single cells were dispensed on the iCell8 platform (Figure 1A). After alignment to the *Drosophila* genome using STAR, downstream analysis and batch effects correction were performed using Seurat ^31^. We filtered cells based on the number of detected genes as well as the percent of mitochondrial genes (Supplementary figure S1A). A total of 2811 hiqh-quality cells from three replicates were used for subsequent cluster analyses. Using hierarchical clustering, we identified a total of 15 cell clusters. Two of these clusters revealed a large overlap of expressed markers and differential gene expression analysis failed to identify markers to discriminate between these clusters. We decided to merge these two clusters (resulting in the cluster “Precursor-PL-1”). For the resulting 14 clusters we identified unique sets of differentially expressed genes (DEG, Figure 1B-C, supplementary data 1-2). All three datasets contributed to each of these 14 clusters (Supplementary figure S1B-D). The name of each cluster corresponds to the name of marker genes or to specific biological features (see subsequent GO term analysis, using DEGs in supplementary data 3-16).

**Figure 1.**
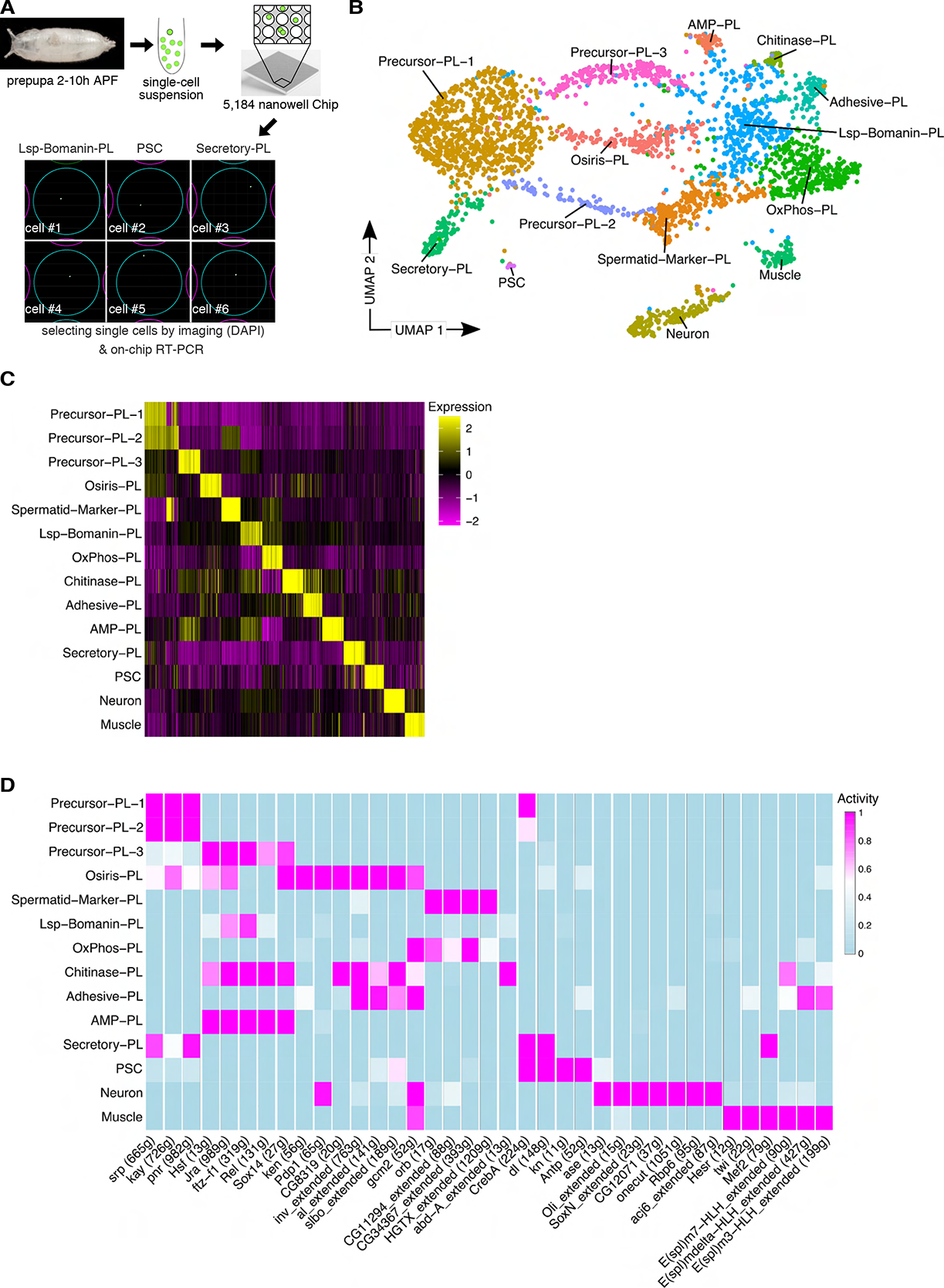
scRNA-seq analysis of pupal *Drosophila* hemocytes reveals a remarkable cellular heterogeneity. A) Workflow for the full-length SMART scRNA-Seq approach using the ICELL8 platform. Hemocytes were isolated from 2-10 h APF *Drosophila* pupae. Cells were stained with Hoechst (additionally, propidium iodide for live cell dataset) prior to their dispensing into the Takara ICELL8 5,184 nanowell chip. Only wells containing single cells were selected using the CellSelect Software prior to the on-chip RT-reaction, library preparation and sequencing. Exemplary images of Lsp-Bomanin-PL, PSC and Secretory-PL cells are shown. B) UMAP plot of 2811 high quality cells in fourteen transcriptomically distinct clusters. C) Heatmap of the top 50 differentially regulated genes per cluster showing transcriptomic differences between cell groups. D) Transcription factor activity of selected transcription factors identified by SCENIC identifies distinct activities between clusters.

To identify gene regulatory networks that characterize the clusters, we performed a regulon analysis using SCENIC ^32^. SCENIC analysis highlights several populations of specific transcription factor activities (Figure 1D, supplementary data 17-18). 12 cell clusters showed significant expression of the pan-hemocyte markers serpent (*srp*) and *hemese* (*he*), but also to a varying extent of known pan-plasmatocyte markers such as the phagocytosis receptor NimC1 (antigen marker P1) and the scavenger receptor Croquemort (Crq) (Supplementary figure S1E). We did not observe a population expressing lamellocyte markers, as expected for non-infested animals (Supplementary figure S1E). Remarkably, we also did not identify a distinct crystal cell cluster in our dataset differentially expressing known transcription factors such as the Runt related transcription factor *lozenge* (*lz*) or *hindsight* (*hnt*) (Supplementary figure S1E). Compared to wandering third instar larvae we only found substantially very few circulating Hnt-positive hemocytes in pupal hemolymph preparations suggesting that crystal cells represent a rare cell type in *Drosophila* pupae and adults as recently reported (data not shown, ^33^).

Two of the clusters contain non-hemocytes that express muscle or neuron specific DEGs and transcription factors, respectively (supplementary figure S2A-D). Muscle- and neuron-specific cell clusters are characterized by increased activities of the transcription factors that specify or maintain muscle or neuronal identity such as *ase*, *onecut*, *Rbp6*, *SoxN*, *Em3-HLH*, *Em7-HLH*, *Emdelta-HLH* and *twi* (Supplementary figure S2E-L).

We previously identified several cytoskeleton and cell motility genes upregulated in pupal hemocytes using a bulk RNA-sequencing approach ^29^. Among the top50 of these genes, 37 genes showed cluster specific expression profiles (Supplementary figure S2M). Many of these genes were highly expressed in Precursor-PL-1, the most abundant cell cluster we identified. This is plausible since bulk RNA-sequencing experiments tend to reflect the expression profile of the most abundant cell types. Other of these genes showed differential expression in OxPhos-PL or Adhesive-PL and Hydrolyzing-PL, suggesting that these populations change transcription profiles with the onset of pupariation to become motile. One gene, *tau*, was specifically detected in a cluster that we later identified as posterior signaling center (PSC) cells. Hemocyte extracts isolated from pupal stages contain the dissolved lymph gland cells, including PSC cells, in contrast to larval hemocyte extracts where the lymph gland is still intact. Thus, the upregulation of tau reflects differences of cell population composition at the onset of pupariation.

### Several plasmatocyte populations serve as precursor cells

Several clusters expressed higher levels of the transcription factor *srp* and SCENIC revealed higher *srp* activity levels in these clusters (Precursor-PL1-3, Osiris-PL and Secretory-PL; figure 2A-B). Expression analysis of a *srp* Gal4 enhancer trap confirmed that *srp* expression is heterogeneous in hemocytes (Figure 2C-D). Previous studies suggested that *srp* expression is higher in undifferentiated hemocytes ^33, 34^. Consistently, cluster Precursor-PL-1 expresses genes involved in hematopoiesis (Figure 2E). We hypothesized that an undifferentiated plasmatocyte population should be present across all stages of hematopoiesis. To test this, we examined the expression of Precursor-PL-1 specific marker genes in datasets containing free larval hemocytes, lymph gland hemocytes as well as in adult hemocytes ^20, 23, 35^. In fact, each of these stages contained a cell population which expressed Precursor-PL-1 markers (PL-0 in larval hemocytes, PM3/4 in lymph gland and Plasmatocytes nAChRalpha3/trol high in adults) (data not shown). This suggests, that a pool of undifferentiated plasmatocytes exists throughout the lifetime of a fly.

**Figure 2.**
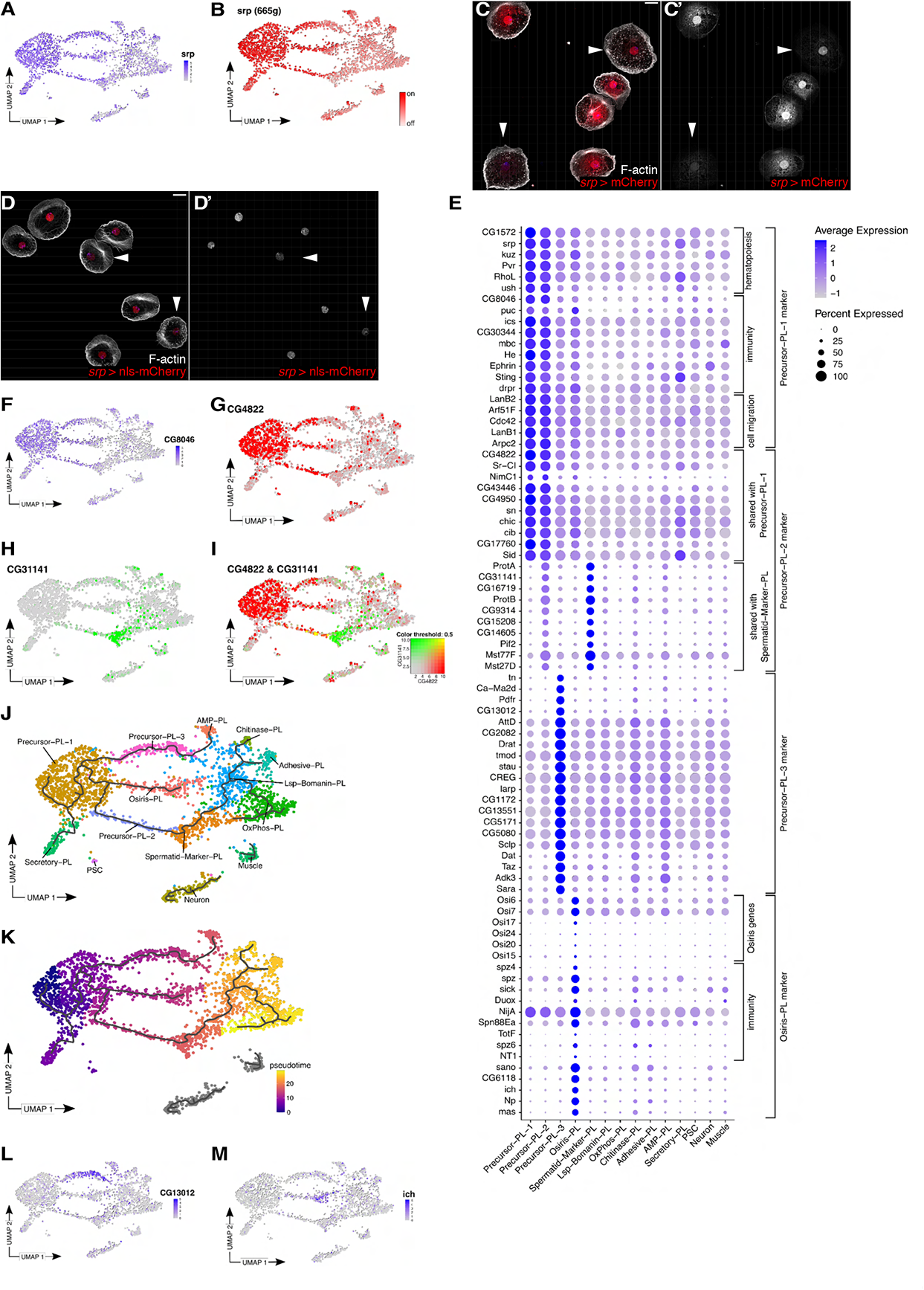
Identification of several precursor plasmatocyte populations. A-B) UMAP plots. *srp* is highly expressed (A) and active (B) in several clusters. C-D) Maximum intensity projection of confocal images of differential expression of the transcription factor *srp* in pupal hemocytes using (C, C’) srpHemo::3xmCherry and (D, D’) srp::Hemo-H2A::3xmCherry; highlighted by white arrowhead are *srp* low-expressing cells; scale bar represents 10 µm. E) Dotplot showing the average expression and percent expression per cluster for marker genes of clusters with high *srp* expression: Precursor-PL-1 – 3 and Osiris-PL. F) Expression of *CG8046*, a Precursor-PL-1 marker, on the UMAP plot. G-I) Expression of Precursor-PL-2 markers on the UMAP plot. G) *CG4822* expression is also detected in Precursor-PL-1. H) *CG31141* expression is also detected in Spermatid-Marker-PL. I) Combined expression of *CG4822* and *CG31141* is only detected in Precursor-PL-2. J-K) Monocle3 pseudotime analysis. J) Pseudotime trajectory on the UMAP plot. K) UMAP plot with pseudotime trajectory and cells colored based on pseudotime. L-M) Expression of marker genes on the UMAP plot. L) *CG13012*, identified as a marker for Precursor-PL-3. M) *ich*, identified as a marker for Osiris-PL.

Markers expressed in Precursor-PL-1 were also detected in other clusters with high *srp* expression (Figure 2F, E). This was particularly true for Precursor-PL-2. However, this cluster differs from Precursor-PL-1 by sharing markers with the low *srp* expressing cluster Glycolytic-PL (Figure 2E, G-I), suggesting that Precursor-PL-2 could be a transition state towards more specified plasmatocyte fate. To test this hypothesis using an unbiased approach, we applied pseudotime analysis using monocle3 ^36^. This method orders cells based on stepwise transcriptomic changes with the most undifferentiated state on one end and more differentiated states on the other end. Indeed, the pseudotime analysis located the Precursor-PL-1 cluster on one end of the trajectory, suggesting that these are the most undifferentiated plasmatocytes (Figure 2J-K). Precursor-PL-2 located between Precursor-PL-1 and Spermatid-Marker-PL, confirming that it is likely a transition state during plasmatocyte differentiation. We also found Precursor-PL-3 to be a transition state towards the more differentiated AMP-PL state. The remaining high *srp* expressing clusters Osiris-PL and Secretory-PL were not identified as a transition state.

Despite overlapping marker genes, Precursor-PL-3 and Osiris-PL can be distinguished from Precursor-PL-1 by the expression of cluster specific markers and cluster specific transcription factor activity as well as the enrichment of distinct GO-terms (Figure 2E, L-M; supplementary figure S3). The same was true for Secretory-PL, which we will revisit later. Osiris-PL expressed many genes important for immunity, providing further support that Osiris-PL are differentiated cells rather than precursor cells (Figure 2E). Additionally, the Osiris-PL cluster is eponymously characterized by the expression of insect-specific gene family Osiris encoding putative transmembrane proteins linked to the development of resistance against a range of different plant and fungi toxins (Figure 2E) ^37–39^. The Osiris-PL cluster showed a characteristic increased activity of *invected* (*inv*), encoding a lineage specific homeodomain transcription factor that is specifically required for anti-fungal defense (Supplementary figure S3B) ^40^. In addition, we identified increased activities of transcription factors selectively controlling a subset of JAK/STAT pathway target genes such as *ken and barbie* ^41^ that in turn regulates *slow border cells* (*slbo*) gene expression (Supplementary figure S3C, D). These data suggest that the PL-Osiris cluster might be a new cell population of specialized hemocytes appearing with the onset of metamorphosis.

### Identification of specialized plasmatocyte subtypes with distinct molecular signatures

We identified six types of *srp* low-expressing hemocyte populations with distinct markers and molecular signatures that indicate that these are specialized immune cells (Figure 3A-H). Many of these clusters were active for *Relish*, a well-known transcription factor that acts downstream of the immune deficiency (IMD) pathway, regulating antibacterial responses (Figure 3L) ^42^. Among them, we identified Spermatid-Marker-PL, Lsp-Bomanin-PL, OxPhos-PL, Chitinase-PL, Adhesive-PL, and AMP-PL.

**Figure 3.**
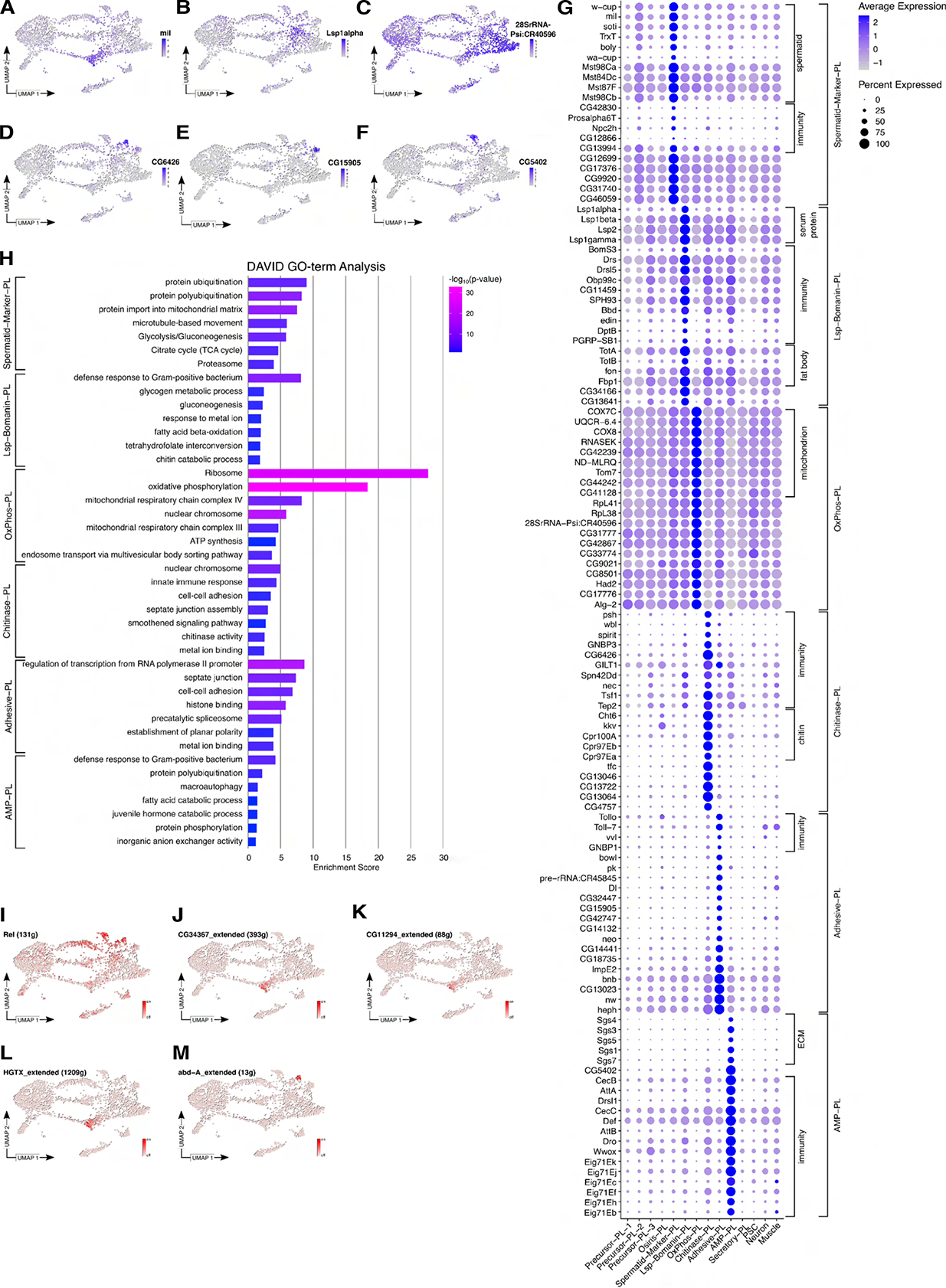
Identification of effector cells with distinct molecular signatures. A-F) Expression of selected markers on the UMAP plot. A) *mil* is a marker for Spermatid-Marker-PL. B) *Lsp1alpha* is expressed in Lsp-Bomanin-PL. C) *28SrRNA-Psi:CR40596* is highly expressed in OxPhos-PL. D) *CG6426* is a marker for Adhesive-PL. E) *CG15905* marks Adhesive-PL. F) *CG5402* marks AMP-PL. G) Dotplot of selected marker genes with average expression and percent expression per cluster for the following clusters: Spermatid-Marker-PL, Lsp-Bomanin-PL, OxPhos-PL, Chitinase-PL, Adhesive-PL and AMP-PL. H) Representative terms of the top seven annotation clusters identified with DAVID 2021 GO-term analysis of differentially expressed genes of the clusters Spermatid-Marker-PL, Lsp-Bomanin-PL, OxPhos-PL, Chitinase-PL, Adhesive-PL and AMP-PL. The barplot height reflects the enrichment score of the annotation cluster and is colored based on the p-value of the GO-term. I-M) Activity of transcription factors on the UMAP plot identified with SCENIC. I) *Rel* is highly active in Precursor-PL-3, AMP-PL and Hydrolyzing-PL. J-L) *CG34367* (J) and *CG11294* (K) and *HGTX* (L) are active in Spermatid-Marker-PL. M) *abd-A* is active in Chitinase-PL.

The Spermatid-Marker-PL cluster expresses immunity genes and a high number of genes that have previously been identified in spermatids (Figure 3A, G). This cluster was further characterized by highly specific activity of several transcription factors including CG34367, CG11294 and HGTX (Figure 3J-L). Among the immunity genes we found a member of the conserved Niemann-Pick type C protein family (Npc2h) protein, which is involved in immune signaling, particularly in the recognition of pathogen related products such as lipopolysaccharides, lipid A, peptidoglycan and lipoteichoic acid ^43^. In addition, Spermatid-Marker-PL cluster is enriched for CG42830 encoding a protein related to Akirins, which play a conserved central role in immune gene expression in insects and mammals, linking the SWI/SNF chromatin-remodeling complex with the transcription factor NF-kappaB ^44^. Furthermore, we identified milkah (mil) as cluster-specific marker encoding a conserved nucleosome assembly factor of the Nap family involved in spermatogenesis and long-term memory formation ^45, 46^. Interestingly, the human homologue, NAP1L has been implicated in stress response and apoptosis through control of the NF-kappaB pathway acting on the anti-apoptotic Mcl-1 gene ^47^. Recent studies identified NAP1L as a novel prognostic biomarker associated with macrophages promoting hepatocellular carcinoma ^48^. High expression of the cytoprotective thioredoxin TrxT against oxidative stress which is also a known conserved marker for inflammatory macrophages further suggests that Spermatid-Marker-PL represents an immune-active plasmatocyte population ^49^(Itoh and Bryson, 2018). Interestingly, a population of Hml positive cells expressing spermatid markers have previously been identified in adult flies and the expression profile of these cells overlapped highly with the pupal Spermatid-Marker-PL cluster. In contrast, we could not identify any clusters among larval or lymph gland hemocytes corresponding to Spermatid-Marker-PL (data not shown).

Lsp-Bomanin-PL expresses several genes encoding larval serum proteins like *Lsp1alpha* and its receptor *Fbp1* (Figure 3B, G). Plasmatocyte populations expressing these genes have previously been identified in larvae ^23, 24^ and consistently, we can identify a larval plasmatocyte cluster (PL-Lsp) with a corresponding expression profile (data not shown). Other genes expressed by Lsp-Bomanin-PL include Bomanin genes like *BomS3* and even secreted anti-fungal and antibacterial peptides such as *Drosomycin* (Drs) or *Drosomycin-like 5* (Drs-5) (Figure 3G). Overall, a GO-term analysis revealed an enrichment for genes important in the defense response to Gram-positive bacteria (Figure 3H). Further, these cells expressed several genes secreted from the fat body including stress-induced humeral factors such as Turandot A and B (TotA, TotB) and the coagulation factor *fondue* (fon) (Figure 3G)^41, 50^.

Thus, the gene expression profile of Lsp-Bomanin-PL partially resembles fat body cells. However, this cell cluster is different from the fat body as it expressed all known hemocyte markers including *srp*, *He*, Pxn and *crq* (Supplementary figure S1E) and its overall gene expression profile correlates well with the other plasmatocyte cell clusters (Supplementary figure S4A). Further, we used ICELL8 imaging data to confirm that the nuclei of these cells were not significantly bigger than other hemocyte populations (27,76 px for Lsp-Bomanin-PL in comparison to 31,97 px for all other plasmatocytes), as one would expect from a fat cell population This suggests a close immune-metabolic interaction between the fat body and *Lsp-Bomanin-PL* cells ^23, 24^.

OxPhos-PL contained mostly cells from dataset2 (Supplementary figure S1C-D) and expressed less genes than other plasmatocyte clusters (Figure S4B). However, the number of detected UMIs as well as the percentage of mitochondrial genes were similar (Fig. S4C-D), leading us to keep this cluster as high-quality cells. OxPhos-PL show a striking increased expression of several genes involved in mitochondrial oxidative phosphorylation system (OXPHOS, figure 3G, H). This cluster also specifically expresses several ribosomal proteins (Figure 3G) which might reflect an important cross-regulatory mechanism between mitochondrial energy production and increased ribosomal assembly and translation as recently described for macrophage tissue invasion in the *Drosophila* embryo ^51^. For professional phagocytes, sufficient ATP as the critical energy source is even required to drive endocytic and phagocytic processes, especially at the onset of metamorphosis when plasmatocytes encounter and clear apoptotic cells from massive larval tissue histolysis. Interestingly, OxPhos-PL plasmatocytes show an increased expression of known actin nucleators and actin-binding proteins required for phagocytosis such as single subunits of the WAVE regulatory complex (WRC; HSPC300), the Arp2/3 complex (Arpc4) suggesting a role of this cluster in the clearance of the larval body during metamorphosis (Figure 3G). OxPhos-PL also strongly expresses the Ribonuclease Kappa (RNASEK) a V-ATPase-associated factor involved in clathrin-mediated endocytosis of large cargo, such as viruses ^52, 53^.

Chitinase-PL show *abd-A* activity and express high levels of immunity relevant genes such as serine proteases like *persephone* (*psh*) and *spirit*, but also diverse regulators of the Toll signaling pathway including *windbeutel* (*wbl*), *Gram-negative bacteria binding protein 3* (*GNBP3*), *Gamma-interferon-inducible lysosomal thiol reductase 1* (*GILT1*), *Transferrin 1* (*Tsf1*) and *Thioester-containing protein 2* (*Tep2*) that mediate the cellular immune response to bacteria (Figure 3G, M). The characteristic high expression of the chitinase *Cht6*, an evolutionarily conserved enzyme involved in ecdysis, organization of the exoskeletal barrier, but also in immune defense in vertebrates, prompted us to name this cluster accordingly (Figure 3G). Recent studies further revealed that the function chitinases are not exclusive to catalyze the hydrolysis of chitin producing pathogens, but also include a crucial role in bacterial infections and inflammatory diseases ^54^. Interestingly our GO-term analysis not only identified the enrichment of genes involved in innate immune response but also an increased expression of genes implicated in cell-cell adhesion and septate junction assembly (Figure 3H), a special feature that we could already observe in our previous bulk RNA seq analysis of pupal hemocytes (Supplementary figure S2M, ^29^. Septate junction (SJ) proteins play an essential role in regulating hematopoiesis in the *Drosophila* lymph gland, but might be also essential for an effective response to parasitic wasp attack ^55 56^. Lamellocytes are known to encapsulate wasp eggs by forming an encapsulation epithelium sealed by SJs that protects the larval tissue ^57^. Hence, Chitinase-PL might assume similar functions.

Similar but even more prominent, the Adhesive-PL cluster expresses several genes implicated in cell-cell adhesion prompting us to name this cluster accordingly (Figure 3G, supplementary figure S4E). The expressed markers range from cadherins like *shotgun* (*shg*), *echinoid* (*ed*) *cad87A*, *fat* (*ft*) to *Fasciclin 3* (*Fas3*), prickle pk and *piopio* (*pio*) known to be involved in epithelial cell polarity. Interestingly, the Adhesive-PL cluster is characterized by low levels of Atilla and Cherrio expression (compared to other clusters, supplementary figure S1E) and might be an intermediate state with the potential to transdifferentiate into terminally differentiated lamellocytes as recently described ^10, 21^

AMP-PL express a high number of immunity genes and our GO-term analysis revealed an enrichment in genes involved in defense to Gram positive bacteria (Figure 3G, H). Among the immunity genes we identified a large repertoire of AMPs like *Attacin-A*, *Attacin-B*, *Drosomycin-like 1*, *Drosomycin-like 6*, *Cecropin A*, *Cecropin B* and *Drosocin* (Figure 3H). In addition, AMP-PL show an increased number of ecdysone inducible genes involved in pupal morphogenesis (*Eig71Ek*, *Eig71Ej*, *Eig71Ec*, *Eig71Ek*, *Eig71Ef*, *Eig71Eh*, *Eig71Eb*) (Figure 3G) ^58^ suggesting that their expression is pupa-specific. Consistently, we did not find any cell types in other developmental stages with a corresponding expression profile suggesting that the AMP-PL is highly determined by the pupal stage (data not shown). Furthermore, AMP-PL produce several extracellular matrix components, including salivary gland secretion proteins (Sgs) which are also induced by ecdysone (Figure 3G) ^59^.

Secretory-PL are among the *srp*-high expressing clusters but without any descending clusters (Figure 2A-B, J-K). This cluster expresses many genes involved in proteolysis with annotated serine-type endopeptidase activity (*CG30098*, *CG31174*, *CG30083*, *CG18636*, *CG14088*) and the intracellular membrane system as well as protein export, inspiring us to name this cluster Secretory-PL (Figure 4A-E, Supplementary figure S5A). SCENIC further identified high activity of cAMP response element binding (CREB) protein *CrebA* a transcription factor regulating components of the secretory pathway ^60^(Figure 4F). In addition, target genes of *dorsal*, the executive transcription factor downstream of Toll signaling pathway were highly expressed in Secretory-PL, confirming the importance of this cell type in innate immunity (Figure 4G). Secretory-PL expressed a highly distinct set of marker genes and shared some marker genes with another cluster that we later identified as PSCs (Figure 4B-E). Shared markers included *Tep4*, *Ance* and *ham* (Figure 4B-D). Using Gal4-enhancer traps and GFP-exon traps for these genes we identified two morphologically distinct cell types: plasmatocyte-shaped cells and small spiky cells (Figure 4H-K). In contrast, *CG31174*, which was not detected in PSCs in our bioinformatic data (Figure 4A) was solely detected in cells with a normal plasmatocyte morphology (Figure. 4K). To confirm that these morphologically different cells are indeed transcriptomic distinct cell types, we co-labeled CG31174 with Tep4 and indeed, while plasmatocyte-shaped cells co-expressed both markers, small spiky cells were never CG31174 positive. Thus, Secretory-PL are plasmatocytes with a classical cellular morphology (Figure 4L). *CG31174* has been recently shown to be expressed in crystal cells ^24^. While we did not detect any classical crystal cell markers in the Secretory-PL cluster (Supplementary figure S1E) we sought to exclude the possibility that these cells are crystal cells. To this end, we made use of datasets available in public single cell databases from various stages ^20, 23, 35^, merged and batch corrected the data with our dataset (Supplementary figure S5B). As suggested by the large shift in gene expression detected in our bulk RNA-sequencing analysis, pupal hemocytes were largely located on a distinct region in the resulting UMAP-plot (Supplementary figure S5C). This confirms that pupal hemocytes are, indeed, transcriptomically distinct from other stages. Due to the availability of cell annotation, we were able to map larval as well as adult crystal cells identified by the authors ^20, 23^. While larval and adult crystal cells located on the same region in the UMAP plot, Secretory-PL located differently, confirming that Secretory-PL are different from crystal cells (Supplementary figure S5D-F).

**Figure 4.**
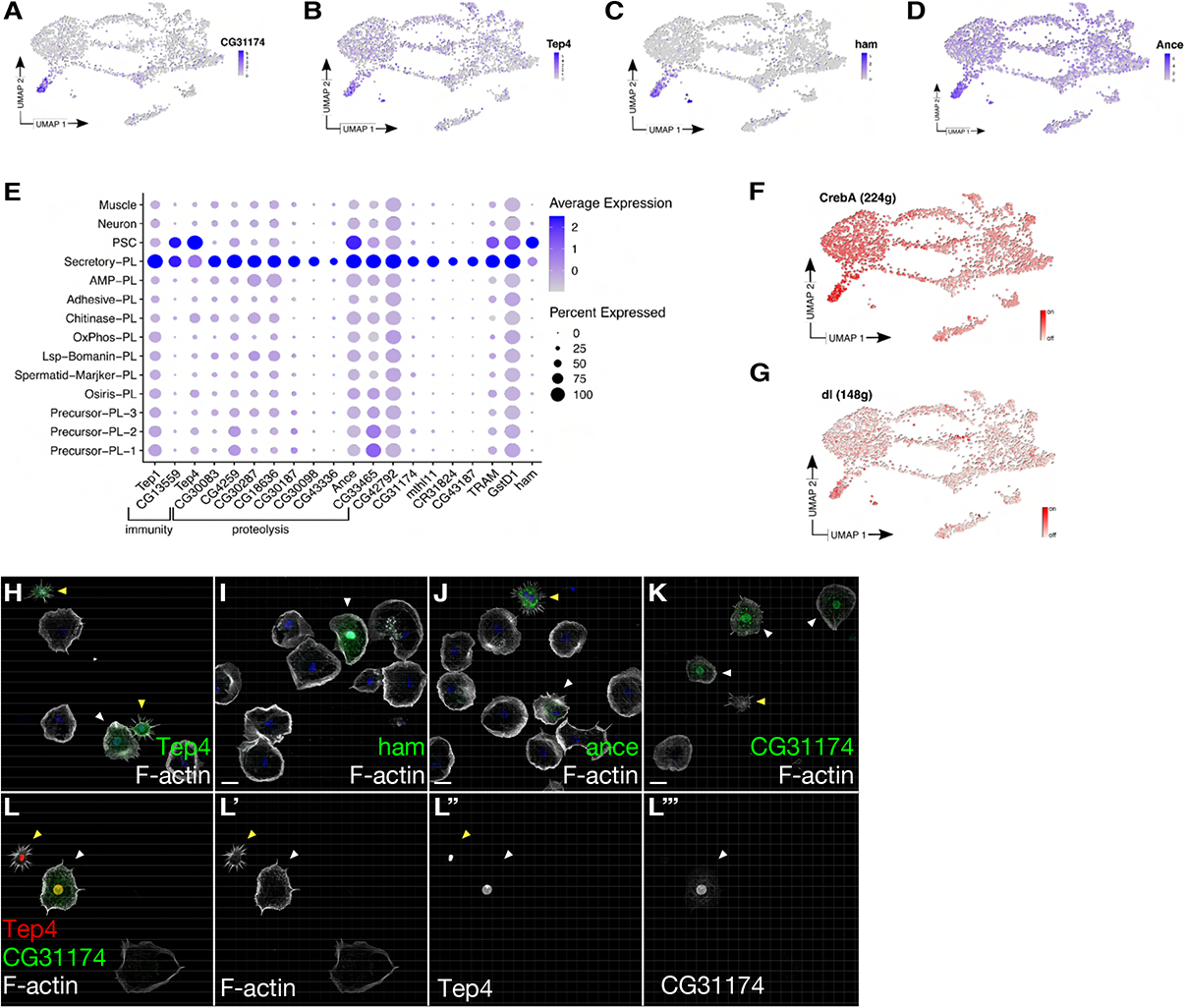
Secretory-PL, a new plasmatocyte type in pupae, is transcriptomically distinct from other developmental stages. A-D) UMAP plots showing the expression of Secretory-PL markers. A) *CG31174* is specific for Secretory-PL. B-D) *Tep4* (B), *ham* (C) and *Ance* (D) expression can also be detected in PSC. E) Average expression and percent expression of Secretory-PL markers on a Dotplot. Note the importance of many genes in immunity or their function in proteolysis. F-G) Activity of transcription factors highly active in Secretory-PL on the UMAP plot as identified by SCENIC. F) *CrebA*. G) *dl*. H-L) Maximum intensity projection of confocal images of pupal hemocytes expressing GFP; plasmatocyte-shaped cells are marked by the white arrowhead; small spiky cells are marked by yellow arrowheads; Alexa568-labeled phalloidin was used to stain the actin cytoskeleton. (H) Tep4-Gal4. (I) Ham-Gal4. (J) Ance ^mimic^-GFP. (K) CG31174 ^mimic^- GFP. (L) Small spiky cells are not CG31174 positive. CG31174 ^mimic^-GFP co-expressing Tep4-Gal4-nls-mCherry. Small spiky cells are marked by a yellow asterisk and plasmatocyte-shaped cells co-expressing both markers are marked by a white asterisk. Scale Bar represents 10 µm

In addition to these major plasmatocyte clusters, our scRNA-seq also identified a small cell cluster that expresses a very unique set of markers, including the early hemocyte marker *srp*, but without detectable levels of more mature hemocyte markers like *He*, *Pxn* and *crq* (Figure 2A, E, S1E, GO-term analysis in S6A). Remarkably, this cell cluster expresses striking markers of the posterior signaling center (PSC) present in the lymph gland, such as the transcription factors *collier* (*kn*) ^61^ *antennapedia* (*antp*) ^62^, but also recently identified new PSC markers such as *tau* ^35^ (Figure 5A-D). Both, *antp* as well as *kn* were also active in the PSC cluster according to SCENIC analysis (Figure 5E-F). We queried datasets containing larval, lymph gland adult hemocytes and found a corresponding cluster in each stage (Supplementary figure S6B-D) ^20, 23, 35^. PSCs originate from the lymph gland and the identification of a PSC cluster in adults suggests that PSCs persist throughout life. The dataset containing larval hemocytes from embryonic origin should, however, not contain lymph gland derived PSCs. The authors suggested that these cells are plasmatocytes with PSC-like expression pattern ^23^. To test whether these cells are indeed distinct from PSCs, we performed cross-stage data integration. We found that pupal and adult PSCs localize at the same region on the UMAP plot with larval PSC-like PL-Impl2 cells (Supplementary figure S6E-G). We then attempted to identify differentially expressed genes across stages of PSC cells, comparing pupal PSCs, adults Hml+ cells with PSC markers, larval PL-Impl2 cells and lymph gland derived cells localizing similarly on the UMAP plot. However, this analysis did not return any significant differences in gene expression among PSC cells from different stages. While this analysis is limited by a low number of cells across all datasets, this result suggests that PSCs remain transcriptionally stable upon lymph gland dissociation. Whether the identification of PSCs in larval stages can be ascribed to issues during dissection or whether some PSCs indeed dissociate from the lymph gland prior to metamorphosis remains to be assessed.

**Figure 5.**
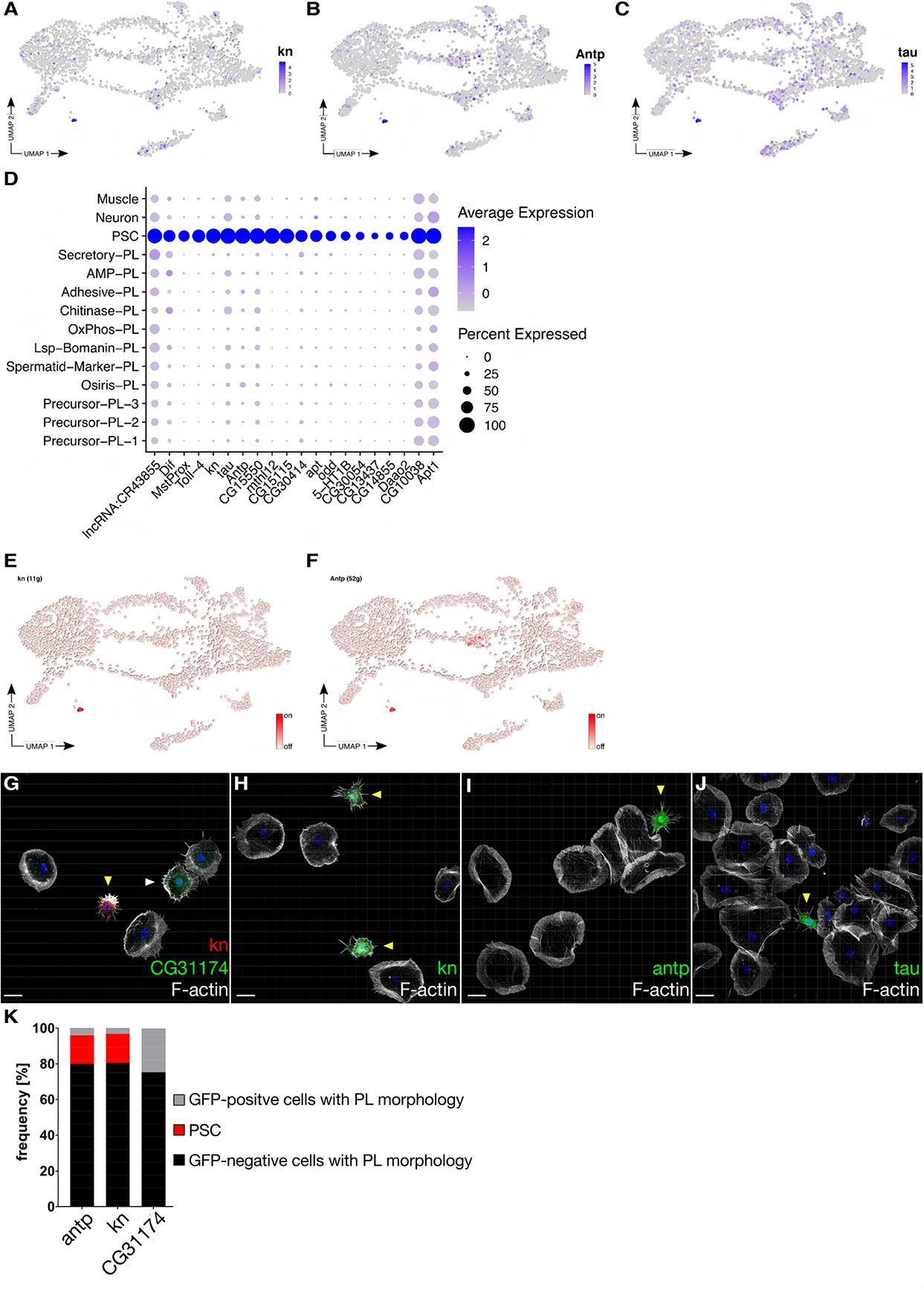
Identification of individual tissue-resident PSC cells in pupae. A-C) Expression of PSC markers on the UMAP plot. A) *kn*. B) *Antp*. C) *tau*. D) Dotplot with average expression and percent of expression of PSC markers. E-F) Activity of transcription factors *kn* (E) and *Antp* (F) on the UMAP plot. G-J) Maximum intensity projection of confocal images of the actin cytoskeleton in pupal macrophages; Alexa Flour labeled phalloidin was used to stain the actin cytoskeleton and DAPI for the nuclei (blue). (G) PSC cells and Secretory-PL cells represent different cell types. CG31174 mimic-GFP co-expressing kn-Gal4-mCherry. PSC cells are marked by a yellow asterisk and CG31174-positive cells are marked by a white asterisk. (H) kn-Gal4 driving UAS-GFP. (I) *antp*-Gal4 driving UAS-GFP. (J) *tau*-Gal4 driving UAS-GFP. (K) Quantification of PSC cells and Secretory-PL cells frequency represented by antp (n=563; GFP+=4%; PSC=16%; GFP-=80%), kn (n=462; GFP+=3%; PSC=16%; GFP- =81%) and CG31174 (n=359; GFP+=24.5%; GFP-=75%) positive cells obtained from six individual experiments. Scale bar represents 10 µm.

We further validated this distinct gene expression profile of PSC cells and characterized them morphologically in more detail. Using transgenic Gal4-enhancer traps and GFP-exon trap fly lines we found that PSC markers *kn*, *antp* and *tau* labeled small spiky cells reminiscent of those cells observed by shared markers (*tep4*, *ham*, *ance*) labelling Secretory-PL (Figure 4H-L, 5G-J). We noticed that *kn* and *antp* occasionally label cells with a more plasmatocyte-like morphology (Figure 5K), in agreement with previous publications and in contrast to *tau*. This population of *kn* or *antp* positive plasmatocytes was less frequent than plasmatocytes labelled by the Secretory-PL marker CG31174, suggesting that they are distinct cell populations (Figure 5K). Indeed, co-staining with CG31174 further confirms that PSC cells are morphologically very distinct from Secretory-PL (Figure 5G).

### PSC cell cluster can differentiate into lamellocytes upon wasp infestion

Different from *kn*, *antp* and *tau*, *Tep4* also marked rare giant cells with lamellocyte morphology in wild type pupae (Figure 6A). Co-staining with an antibody against the lamellocyte marker gene Atilla further confirmed Tep4-positive terminal differentiated lamellocytes (Figure 6B). Remarkably, wasp infestion dramatically increased the number of Tep4-positive lamellocytes (Figure 6C, quantification in Figure 6D) suggesting that Tep4 marks a group of plastic cells which are capable of differentiation into lamellocytes. Upon wasp infestion, induced lamellocytes could also be marked by *kn* expression in pupae (Figure 6E). To test whether these lamellocyte descend from PSCs, we made use of the G-TRACE (GAL4 Technique for Real-time And Clonal Expression) lineage tracing system (Figure 6F) ^63^. This method uses a cell type specific Gal4 driver and a UAS-RFP construct to mark Gal4 positive cells with RFP. Gal4 also induces Flp expression, which removes the stop codons in a ubi-FRT-Stop-FRT-GFP construct, resulting in stable GFP expression. Hence, Gal4 positive cells are RFP and GFP positive, while offspring cells are GFP positive but RFP negative. Remarkably, G-TRACE lineage analysis using *kn*-Gal4 resulted in the detection of positive lineage traced (GFP-positive) cells (Figure 6G-G’’’). However, past studies using *kn-Gal4* induced G-TRACE have identified a population of low *kn* expressing plasmatocytes that do not fully differentiate and may serve as an adult hematopoietic niche along the body wall of the fly abdomen ^19, 33^. Indeed, while *kn* > GFP cells correspond to the small and spiky PSC morphology, a small number of GFP positive cells displayed plasmatocyte typical morphology (Figure 5K). Hence, to test whether low *kn* plasmatocytes or high *kn* expressing PSCs are precursors of lamellocytes, we combined G-TRACE with *tau-Gal4*, which is specific for PSCs. Indeed, when we wasp infested larvae with *tau* driving G-TRACE, we solely observed large GFP positive cells with a morphology typical for lamellocytes (Figure 6H-H’’’). This confirms that the cells of the PSC cluster are able to differentiate into lamellocytes upon parasite infection and suggests that the PSC apart from being a lymph gland hematopoietic niche also functions as a cell reservoir to pupal and adult blood cells.

**Figure 6.**
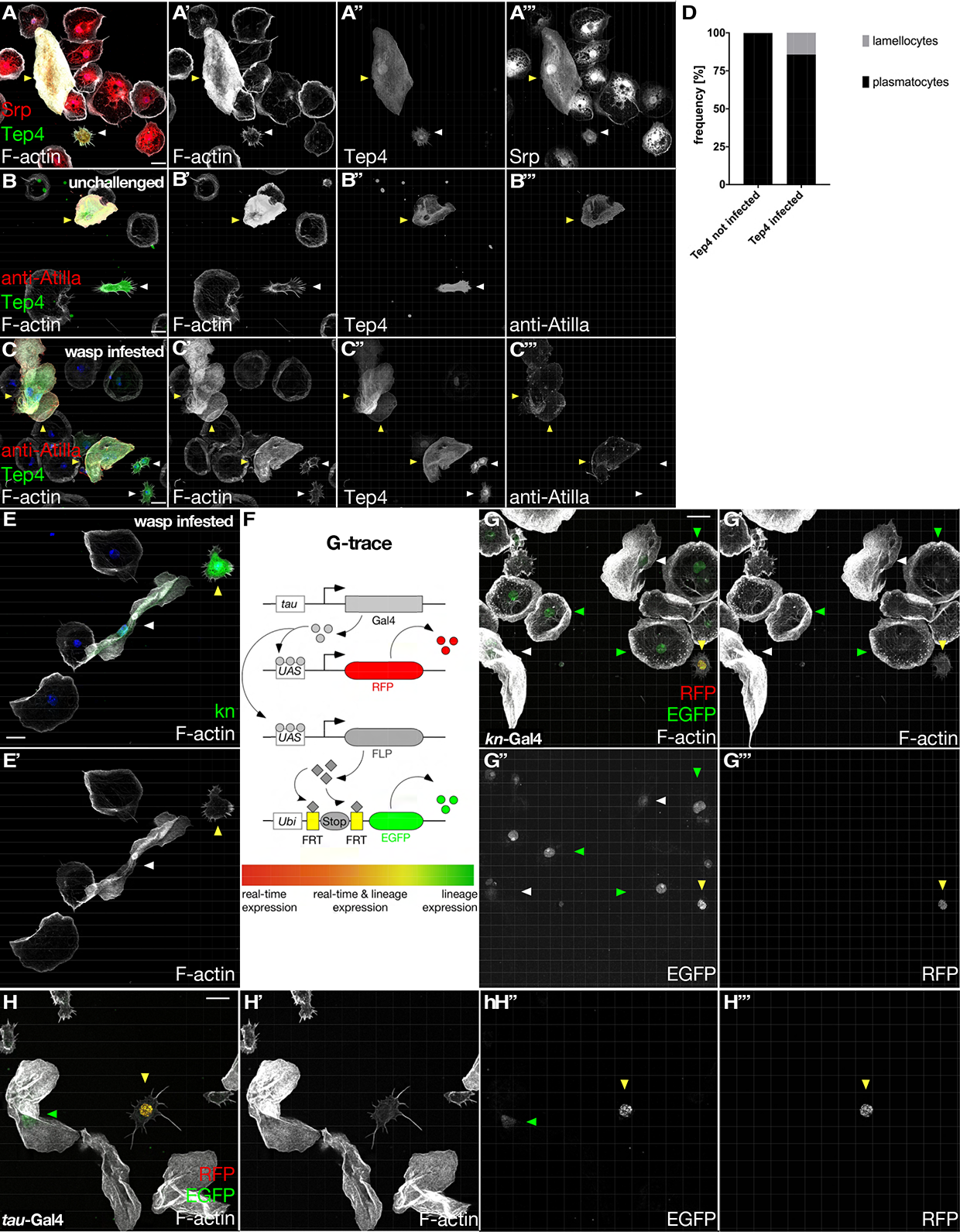
PSC cells can differentiate into lamellocytes upon wasp infestion. A-C) Maximum intensity projection of confocal images of pupal macrophages; Alexa568-labeled phalloidin was used to stain the actin cytoskeleton. (A) Co-expression of srpHemo::3xmCherry and Tep4-Gal4 > UAS-GFP. Spiky small cells express the transcription factor *srp*. Tep4 and *srp* double positive cells are marked by white arrowheads. (B, C) Tep4-Gal4 > UAS-GFP cells were stained for the lamellocyte marker gene atilla (red) either in unchallenged condition (B) or upon wasp infestion (C). Large Tep4-expressing cells show characteristic lamellocyte morphology, marked by a yellow asterisk. (d) Quantification of Tep4 expressing lamellocytes enrichment upon wasp infestion. Infected n=429 cells and control n= 443 cells obtained from three individual experiments. (e) kn-Gal4 > UAS-GFP individuals were infested with wasps. Upon infestion GFP positive cells are either small and spiky (yellow arrowhead) or large with lamellocyte typical morphology (yellow arrowhead). f) Schematic overview of the G-TRACE system. Gal4 protein is specifically expressed in PSCs under the control of the *tau* promoter. PSCs are RFP positive due to the UAS-RFP transgene. Gal4-positive PSCs induce flipase (FLP) expression via UAS-FLP. FLP can remove an FRT-flanked stop codon downstream the ubiquitous ubi promoter. Stop codon removal leads to stable and inheritable expression of GFP, which allows tracing of offspring cells produced by PSCs. Differentiating PSC offspring turn off *tau-*Gal4 resulting into GFP positive but RFP negative cells. (G, H) Cell lineage analysis of pupal hemocytes using (G) *kn* > G-TRACE. PSCs are small and spiky cells double positive for RFP and GFP (yellow arrowhead). *kn* > GFP cells correspond to the small and spiky PSC morphology, a small number of GFP positive cells displayed plasmatocyte typical morphology (green arrowheads), GFP positive lamellocytes are marked by white arrowheads. (H) *tau* > G-TRACE. Some lamellocytes are GFP positive (green arrowhead) showing that they derived from PSC cells. Other lamellocytes are GFP negative and may thus differentiated from other cellular sources. Scale bar represents 10 µm.

### PSC cells are migratory immune responsive cells persisting into adult

To better characterize PSC cells *in vivo*, we used the *kn*-Gal4 driver to perform live-cell imaging in developing pupae (Figure 7). While kn-Gal4 also occasionally marks plasmatocytes (Figure 4K), our in-depth analysis of different PSC markers allowed us to determine that PSCs are smaller with a distinct morphological shape. We therefore focused our analysis on cells with the typical PSC morphology. As previously reported ^64–66^, prominent *kn* expression is initially found in dorsal muscle fields in the thoracic and abdominal segments of early pupae (Figure 7A), and its expression becomes restricted to segmental abdominal muscles, along the anterior-posterior border of the developing wing and in pericardial cells along the dorsal vessel in later pupal development (>16h APF-96h APF; Figure 7B, C). High-resolution spinning disc live cell imaging microscopy of 4h APF old pupae revealed migrating single *kn*-marked PSC cells that form polarized dynamic lamellipodial protrusions and start to redistribute from the dorsal patches of the body wall as similarly observed for plasmatocytes (Figure 7D; Supplementary movie M1) ^9, 29^. *kn*-marked PSC cells are highly motile but much smaller as compared to *hml*-marked plasmatocytes, confirming that we indeed observed PSCs when focusing on small *kn* positive cells instead of plasmatocytes (Supplementary movies M1 and M2). This difference in cells size becomes more evident in later pupal development (> 16h APF) when cell dispersal in the pupa is further advanced and cell numbers are increased (Figure 7E, F). Compared to *hml*-marked plasmatocytes, which are more evenly distributed in the whole developing pupa, *kn*-marked PSC cells become more restricted to the abdomen during pupal development until hatching (compare stages 16h APF and 96h APF in Figure 7 B, B’ and C, C’). Remarkably, PCS cells are not only highly motile, but also able to phagocytose bacteria although with limited capacity compared to *hml*-marked plasmatocytes (Figure 7G). Finally, we tested whether PSC are immune-responsive cells using laser-ablation experiments (Figure 7H, supplementary movie M3. Upon wounding PSC cells switch from random to directed migration towards the wounding site as indicated (Figure 7H). No significant differences in directionality were found between PSC cells and *hml*-marked plasmatocytes as quantified by measuring the cell bias angle (Figure 7I). Combined, these data strongly suggest a dual function of PSC cells in *Drosophila* hematopoiesis. Apart from its well-known important function as larval hematopoietic progenitor niche, PSC cells can function as tissue resident immune-responsive cell reservoir with limited phagocytic ability but still capable of blood cell differentiation upon immune challenge later in development.

**Figure 7.**
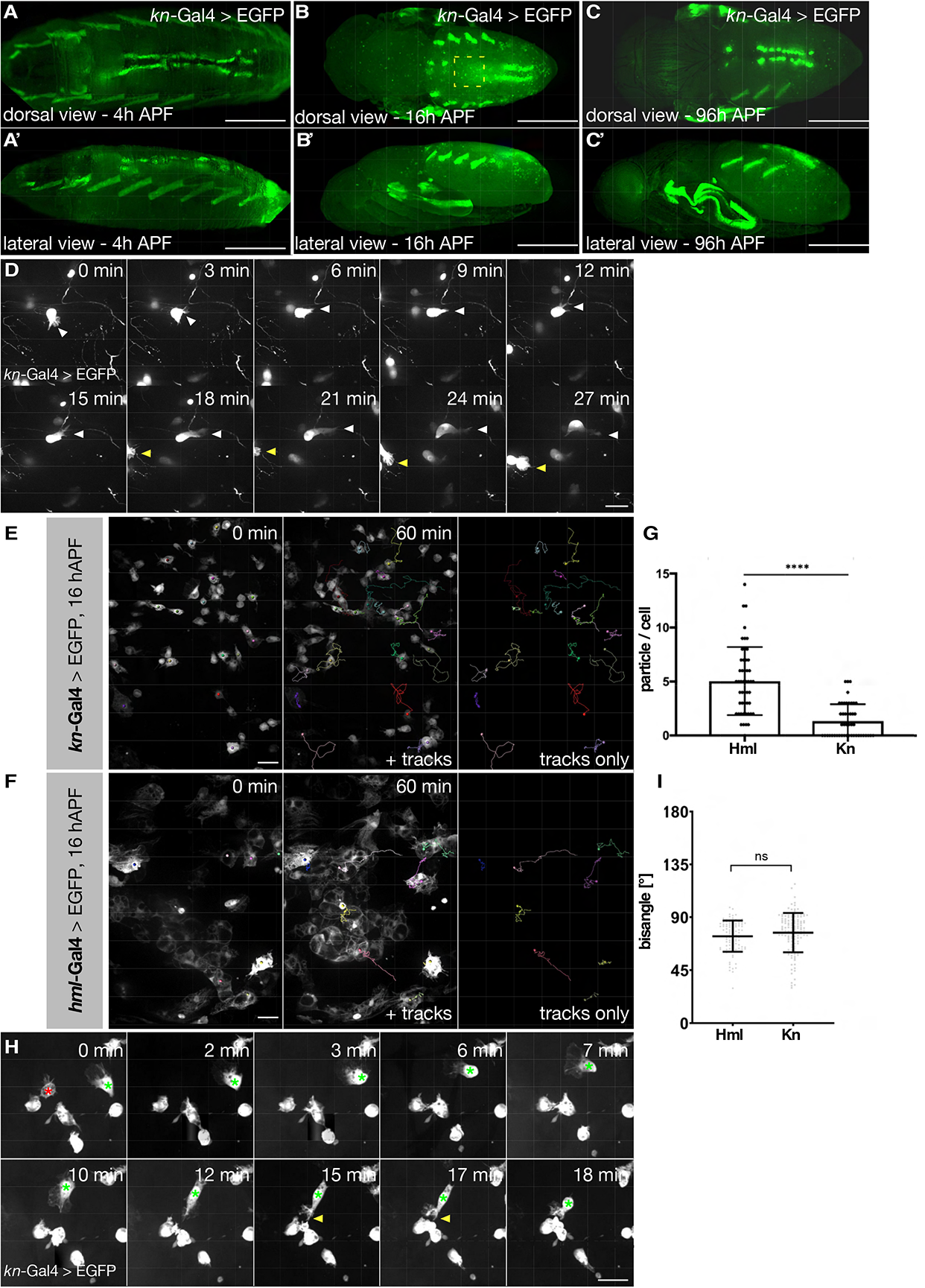
PSC cells are motile tissue-resident immune responsive cells persisting into adult. Representative *in vivo* images of GFP expression under the control of the collier promotor *kn*-Gal4 (PSC cells are marked). Scale bar represents 50 µm. (A) dorsal and (Á) lateral view of a pupae 4h APF. (B) Dorsal and (B’) lateral view of a pupae 16h APF. The rectangle marks the region of accumulation of PSCs visualized by spinning disc microscopy. The expression on the pupal wing represents the A/P boundary. (C) Dorsal and (Ć) lateral view of a pupae 96h APF. (D) Frames of a spinning disc time-lapse movie of randomly migrating 4h APF *kn* > GFP pupal hemocytes. Images were taken every 20 s for 30 min. A white and a yellow arrowhead mark the migrating PSC hemocytes. The time point of each image is annotated, scale bar is 20 µm. (E, F) Still images of spinning disc time-lapse movies of directed macrophage migration upon wounding of a single cell in the *kn*-Gal4, UAS-GFP (E) or *hml*-Gal4, UAS-GFP (F) background. Cells were imaged for 60 min after laser ablation in 30 s intervals and tracked by Imaris software. (E) Trajectories indicate that GFP expressing PSC hemocytes are highly motile. (F) Hml-marked plasmatocytes are much bigger and contain more phagocytic vacuoles than PSC cells. (G) Quantification of phagocytic activity of *hml*-positive plasmatocytes and PSC cells marked by *kn* expression measured by the uptake of Texas-Red-conjugated *E. coli* particles per cell. PSC cells are phagocytotic but show a significantly reduced phagocytic activity compared to cells marked by *hml*-GFP expression. *hml* (n= 50), *kn* (n= 50) positive cells were used from 4 (*hml*) and 5 (*kn*) individual experiments, respectively. *P-values* from one-way ANOVA and Tukeýs multiple comparisons tests are shown. (\*\*\*\**P<*0.0001*; ***P*=0.0002; \*\**P*=0.0023). (H) Magnification of PSC macrophages close to the ablated cell (marked by a red asterisk) at the indicated time points. Green asterisk marks a migrating wound-responsive PSC hemocyte towards the ablation site. (I) Quantification of the directionality of cells marked with *hml* or *kn* towards the wound. Directionality is described with the bias angle of the migrating cells. *kn* positive PSC cells show a similar wound response as *hml-*GFP cells, which serve as a pan-hemocyte control. *kn* (n=129) *hml* (n=103) values from 10 individual experiments. Graphs are depicted in a scatter dot blot with bars indicating mean and standard deviation. ns = p = > 0.03 (Welch’s t test).

## Discussion

In this study we present a comprehensive single-cell transcriptome analysis of *Drosophila* pupal hemocytes at the onset of metamorphosis when most larval structures are degraded and replaced by adult tissues and organs. This developmental stage represents a striking example of dramatic and systemic physiological changes that requires an integration of the innate immune system. In response to ecdysone, hemocytes rapidly upregulate cell motility and phagocytosis of apoptotic debris, and acquire the ability to chemotax to tissue damage. ^9, 27, 28, 67^. Recent bulk RNA-seq analysis revealed 1542 differentially regulated genes, from which 804 genes are up-regulated in pupal hemocytes compared to quiescent larval hemocytes ^29^. These striking differences in the global gene expression likely reflect both transcriptional changes of embryo-derived hemocytes and the emergence of new progenitor-derived hemocytes of the lymph gland since both hemocyte lineages mix together and disperse in the pupa ^68^. Consistently, our single cell RNA-seq data analysis revealed both cell clusters common in other stages of hematopoiesis and new subgroups appearing with the onset of metamorphosis and persisting into adulthood. The most abundant *srp*-high expressing Precursor-PL-1 represents a good example for a cluster present across all stages of hematopoiesis, annotated as PL-0 in larvae ^23^, as PM3/PM4 in lymph glands ^35^ and as nAChRalpha3/troll-high in adult flies ^20^. Precursor-PL-1 contains undifferentiated cells with a broad gene expression profile and no distinctive markers; thus, they might function as large plasmatotcyte reservoir throughout the entire life cycle.

In comparison to that, Lsp-Bomanin-PL only correlates with the comparable PL-Lsp cluster in larvae, however, no correlations are found in lymph glands or adult suggesting that this cell population derives from the embryonic/larval lineage with distinct functions in metamorphosis. By contrast, Spermatid-Marker-PL only has a common expression profile with the *Hml+ with sperm markers* cluster found in adults ^20, 23, 35^ suggesting that it represents an immune-active plasmatocyte population with important functions beyond metamorphosis.

Remarkably, most of the plasmatocyte clusters with distinct molecular signatures are only found in the pupa without significant correlations with cells from different developmental stages that highlight the diverse hemocyte function during pupal development. The Chitinase-PL and Adhesive-PL clusters might represent such effector cells both sharing an increased expression of several genes implicated in cell-cell adhesion and septate-junction formation required for encapsulation of parasitic wasp egg. Most known parasitoid wasp species attack the larval or pupal stages of *Drosophila*. While *Trichopria* drosophilae infect the pupal stages of the host, females of the genus *Leptopilina* and *Ganaspis* attack the larval stages ^69, 70^. Core components of Toll pathway are highly upregulated in Adhesive-PL that control the immuno-genetic circuit of host immune response against parasitoid wasp attack by activating the NF-kappaB transcription factor, *Dorsal*. Future high-resolution live-imaging combined with genetics will be further required to dissect in more detail how these newly identified plasmatocyte effectors contribute in this fascinating process of encapsulation.

Many of the remaining plasmatocyte populations express pupal specific expression profiles suggesting a specification for tissue clearance, a conserved physiologically important process during development and tissue remodeling ^71^. Thus, these hemocyte populations and their expression profiles may therefore be of particular interest for future studies in this context.

Besides the identification of new specialized plasmatocyte subpopulations that first appear during pupal development, we identified a small group of immune cells that resembles the PSC, a well-known stem cell niche that controls the differentiation of effector types from progenitors in the lymph gland, but also participate in the larval response to wasp parasitization ^72^. PSC cells are clearly distinguished by their co-expression of Antp and Collier/knot from other regions within the lymph gland ^62, 65, 73^. Previous work has shown that PSC cells reside in the lymph gland and provide signals to regulate progenitor maintenance or differentiation ^17, 62, 65, 73^. In this work, however, we identified PSC cells from the hemolymph of early pupa that are characterized by the expression of Antp and Collier (Kn), but also by the recently identified new PSC markers such as Tau ^35^. High-resolution live-imaging microscopy analysis using a *kn*-Gal4 enhancer trap further confirmed the existence of single motile and immune-responsive PSC cells persisting throughout pupal development, thus, far longer than described for progenitor cells from histolyzing lymph glands ^74^.

Our results demonstrate that both *kn*-but also *tau*-traced progenitors are able to differentiate into lamellocytes in response to wasp infection. While *kn* positive cells contain PSCs as well as other immune cell populations ^19, 33, 75^, the identification of *tau* as a specific marker for PSCs allowed us to determine the PSCs as the cell pool transdifferentiating into lamellocytes. This suggests that PSC cells act not only as a stem cell niche in larval hematopoiesis, but can also contribute as cell reservoir to pupal and adult blood cells. Recent studies have identified several types of niche cells, including niches in the *Drosophila* reproductive organs as well as rat limbal niche cells, with the potential to transdifferentiate into cells of the lineage they normally regulate ^76–78, 79, 80^. Considering the growing body of literature attending to transdifferentiation upon environmental stressors including in cells of the human immune system ^78, 81–83, 84–86^. This study together with others emphasizes the importance of niche cells in this process.

Although our G-TRACE lineage analysis reveals that PSC cells can transdifferentiate in response to an immune challenge, and cells persist to adults ^19, 33^, it still remains unknown whether PSC cells substantially proliferate providing an active hematopoietic hub in *Drosophila* adults as previously suggested for progenitor cells from the tertiary and quaternary lobes of the larval lymph gland ^33^. Recent findings contradicted an adult hematopoiesis model, but rather provided evidence that mainly embryo-derived hemocytes relocate to the respiratory epithelia and the heart where they provide a major reservoir to mediate a protective humoral immune response in the adult fly ^67^. These embryo-derived reservoir hemocytes, however, were characterized by the plasmatocyte-specific *Hemolectin* (*Hml*) Gal4 driver. PSC cells are not marked by *hml-*Gal4, but rather co-express *srp* suggesting the existence of different reservoirs derived from different pools.

In summary, this study identified new pupal precursor and effector hemocytes with distinct molecular signatures and cellular functions clearly distinct from other stages of hematopoiesis. Our data further uncovers an additional route for lamellocyte differentiation. Besides the transdifferentiation from existing plasmatocytes or from differentiating lymph gland prohemocytes, we provide first evidence that PCS cells can differentiate into lamellocytes. This work will allow future genetic studies to better understand immune cell plasticity and function during metamorphosis and innate immunity.

## Experimental procedures

### Drosophila Genetics

Fly husbandry and crossing were carried out according to the standard methods. Crosses were maintained at 25°C. The following fly lines were used: *hml*-DsRed (D. Siekhaus), srpHemo-3xmCherry ^87^ *hml*Δ-Gal4, UAS-eGFP ^88^; UAS-eGFP (BL 6874). tep4-Gal4 (BL 76750), Ance^MiMiC^-GFP (BL 59828), kn-Gal4 (BL 67516), pcol85-Gal4 ^65^, Antp-Gal4 (BL 26817), ham(GMR80G10)-Gal4 (BL 40090), tau-Gal4 (BL 77641), CG31174^MiMic^-GFP (BL 24040); UAS-mCherry-NLS (BL 38425). G-TRACE (UAS-RedStinger, UAS-FLP, Ubi-p63E(FRT.STOP)Stinger) (BL 28281). For wasp infection, flies were transferred into a fresh food vial and placed at 25°C for 48h. Flies were then removed and larvae submitted to egg laying by wasps of the species *Leptopilina boulardi,* strain G486. Wasps were removed before dissecting of pupae as described later.

### Isolation of Drosophila hemocytes for single-cell RNA-sequencing

50-60 2-10h APF pupae were collected and washed in 1 x PBS. The pupae were then transferred to 1 x Schneideŕs *Drosophila* medium (Gibco) supplemented with 10% FBS, 50 units/mL penicillin, and 50 µg/mL streptomycin and opened using forks to rinse out the hemolymph. Cells were filtered through a 50-µm cell strainer and collected in a total volume of 500 µL. Subsequently, centrifugation at 500 x g and 4°C for 20 minutes was performed. The supernatant was discarded, and the pellet was resuspended with 500 µL PBS + 0.01 % BSA for single-cell sequencing of live cells. For fixed samples, the previous pellet was resuspended with 100 µL 1 x PBS and subsequently fixed by adding 4 volumes of ice-cold 100% methanol (final concentration of 80% methanol in PBS) and thoroughly mixed with a pipette. Cells were stored at −20°C until use (1-3 days). For single-cell sequencing, cells were moved to 4°C and kept on ice throughout the procedure. Fixed cells were pelleted at 500 x g for 15 minutes and rehydrated in 500 µL PBS + 0.01 % BSA. For datasets 1 and 2 we used fixed samples, while we used live cells for dataset 3.

### Single-cell RNA-sequencing of Drosophila pupal hemocytes

The Takara ICELL8 5,184 nano-well chip was used with the full-length SMART-Seq ICELL8 Reagent Kit. Cell suspensions were fluorescent-labelled with live/dead stain, Hoechst and propidium iodide (NucBlue Cell Stain Reagent, Thermo Fisher Scientific) for 15 min prior to their dispensing into the Takara ICELL8 5,184 nano-well chip. CellSelect Software (Takara Bio) was used to visualize and select wells containing single and live cells. Next, cDNA was synthesized via oligo-dT priming in a one-step RT-PCR reaction. P5 indexing primers for subsequent library preparation were dispensed into all wells receiving a different index, in addition to Terra polymerase and reaction buffer. Transposase enzyme and reaction buffer (Tn5 mixture) were dispensed to selected wells. P7 indexing primers were dispensed to wells. Final Illumina libraries were amplified and pooled as they were extracted from the chip. Pooled libraries were purified and size selected using Agencourt AMPure XP magnetic beads (Beckman Coulter) to obtain an average library size of 500 bp. A typical yield for a library comprised of ∼ 1,300 cells was ∼ 15 nM. Libraries were sequenced on the HiSeq 4000 (Illumina) to obtain on average ∼ 0.3 Mio reads per cell (SE; 50 bp). A list of Gal4-enhancer trap and GFP-exon trap fly lines used for ex vivo validation can be found in Supplementary data 19.

### Bioinformatic analysis

Raw sequencing files (bcl) were converted into a single fastq file using Illumina bcl2fastq software (v2.20.0.422) for each method. Each fastq file was de-multiplexed and analyzed using Cogent NGS analysis pipeline (CogentAP) from Takara Bio (v1.0). In brief, “cogent demux” wrapper function was used to allocate the reads to the cells based on the cell barcodes provided in the well-list files. Subsequently, “cogent analyze” wrapper function performed a preliminary analysis, including: read trimming with cutadapt (version 3.2); genome alignment to Drosophila M. Version 103 using STAR (version 2.7.2a); read counting for exonic, genomic and mitochondrial regions in Drosophila M. genes from ENSEMBL gene annotation version 103 using featureCounts (version 2.0.1); and summarizing the gene counts into gene matrices with number of reads expressed for each cell in each gene. Raw gene matrices underwent quality-control filtering for cells and genes using the following parameters: for cells, only those with at least 10,000 reads associated to at least 300 different genes were kept, and for genes, only those containing at least 100 reads mapped to them from at least 3 different cells were kept. Subsequent analyses were performed using Seurat v4.1.1 in Rstudio 2022.02.2. Low-quality cells were filtered out based on the number of detected genes and percentage of mitochondrial genes (Supplementary figure S1A). Batch correction between the three replicates was performed, UMAP coordinates were calculated and unsupervised clustering was perfomed with 16 dimensions and the default resolution factor. Clustering was controlled by testing for sufficient distinct marker genes using the FindAllMarkers() command and two clusters were merged due to insufficient distinct markers (resulting in Precursor-PL-1) (Script 1). Pseudotime analysis was performed using monocle3 v1.0.0 using UMAP coordinates calculated with Seurat v4.1.1 and with learn_graph_control = list(ncenter=480) in the learn_graph() command (Script 2). SCENIC analysis was performed with SCENIC v1.3.1 using the cisTarget v8 motif collection mc8nr (Script 3). For a GO-term analysis we used differentially expressed genes identified using FindMarkers(). Gene names were converted to FlybaseIDs and analysed with DAVID 2021. Cross-dataset analysis was performed using Seurat v4.1.1 including batch correction and UMAP calculation (Script 4).

### Data and Code availability

Raw data will be submitted to GEO and made publicly available upon manuscript acceptance. Output files as well as data associated with figures are published as Supplementary data files. In addition, the final processed data will be made available for download from Mendeley as a .rds file. To allow easy use of this file, we provide Script 5, which aims to guide users without prior knowledge in R. Code used in this study is submitted in Scripts 1-4 and any further details can be obtained upon request.

### Immunohistochemistry

Pupal macrophages were isolated as described previously ^89^. In short, white to light brownish prepupae (2-4h APF) were collected and washed in 1 x PBS. The prepupae were then transferred to 1 x Schneideŕs *Drosophila* medium (Gibco) supplemented with 10% FBS, 50 units/mL penicillin, and 50 µg/mL streptomycin and opened using forks to rinse out the hemolymph. Cells were spread on glass coverslips, previously coated for 30 minutes with ConcanavalinA (0.5 mg/mL, Sigma), for 1 h at 25°C. The supernatant was removed and adherend cells were subsequently fixed for 15 min with 4% paraformaldehyde in 1 x PBS at RT. Cells were treated shortly with 1 x PBS + 0.1 % Triton X-100 followed by three washing steps with 1 x PBS. If no antibody staining was performed, the treatment with PBS-T was omitted. Cells were stained with antibody for 2 h at RT and secondary antibody with Phalloidin and DAPI for 1 h at RT in a humidified dark chamber. Stained cells were mounted in Mowiol 4-88 (Carl Roth). The following antibodies were used: anti-Atilla (1:10), anti-Hnt (1:5, 1G9 from DSHB). The following secondary antibody were used: polyclonal Alexa Flour-647-conjugated goat-anti-mouse (1:1000 dilution; #A21236, Invitrogen). F-actin staining was visualized using Phalloidin-iFlour 405 (1:100 dilution, Abcam #176752) or Alexa Flour-Phalloidin 568 (1:100 dilution, #A12380, Invitrogen) and nucleus by DAPI staining (1 µg/mL, #62248, Thermo Scientific).

### Phagocytosis assay

For phagocytosis assays, pupal macrophages were isolated the same way as for immunohistochemistry. For phagocytosis assays, 1nM phenyl thiourea was added to the preparation medium. After adhering to a glass coverslip for 1 h at 25°C, the medium was gently removed, and replaced with 200µl of fresh medium supplemented with 1nM phenyl thiourea and Texas red-conjugated *Escherichia coli* K-12 particles (Molecular Probes BioParticles® E-2863) for a density of 53000 particles/mm^2^ and incubated for 1 h at room temperature in the dark. Cells were then fixed by incubation in 4% paraformaldehyde solution for 15 minutes at RT. The number of internalized particles per cell was counted for 50 cells per group from confocal fluorescent imagers taken Leica TCS SP8 with an HC PL APO CS2 63x/1.4 oil objective and Leica Application Suite X (LasX, Version 3.5.2.18963). The results were analyzed by one-way ANOVA and Tukey’s multiple comparisons test in GraphPad Prism version 7 by GraphPad Software.

### Image acquisition and microscopy

Pupae were imaged as a whole with Leica Fluorescence Stereo Microscope and Leica Application Suite X (LasX, Version 3.5.2.18963).

Confocal fluorescent images were taken with a Leica TCS SP8 with an HC PL APO CS2 63x/1.4 oil objective and Leica Application Suite X (LasX, Version 3.5.2.18963). Live imaging of macrophage cultures was performed using a Zeiss CellObserver Z.1 with a Yokogawa CSU-X1 spinning disc scanning unit and an Axiocam MRm CCD camera (6.45 µm x 6.45 µm) and ZenBlue 2.5 software. Ablation experiments were done using the UV ablation system DL-355/14 from Rapp OptoElectronics, as reported previously ^9, 89^.

### Live-cell imaging of pupal macrophages

Live imaging of pupal macrophages 4 h APF for random migration as well as 16 h APF for directed migration experiments were performed as described previously ^89^. 4 h APF prepupae were collected and glued to a glass coverslip on their dorsal-lateral side. Spinning disk time-lapse movies were taken with images every 20 s for 30 min. 16-18 h APF pupae were dissected out of their cuticula and laid on a glass coverslip on their dorsal abdomen. Single cells were ablated as described in the previous chapter. Spinning disk-time-lapse movies were taken every 30 s for 60 min.

### Quantification of directed migration of macrophages

Tracking of migrating macrophages was performed using the spots module in the Imaris 9.3 (Bitplane) software. The reference frame module was set at the ablation site. After automatic tracking, all time laps movies were checked for correct results and were manually corrected. The bias angle between the vector toward the ablated cell and the direction vector of the cell was calculated in R (R Studios Version 1.4). The angle between the vector directly towards the ablation cell and the direction vector of the cell at each time point was calculated with 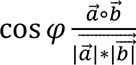,

i.e., the scalar product 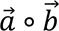 of vectors *a* and *b* divided by the multiplication of the length of each vector 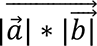.

For directed migration only cells within a 10-80 µm radius of the wounding site were analyzed. Results were statistically analyzed using GraphPad Prism 7 by GraphPad Software.

## Acknowledgements

We thank the Bloomington Stock Center and VDRC for fly stocks. We thank Daria Siekhaus for sharing fly stocks. The work was supported by grants to S.B. (BO1890/5-1) from the Deutsche Forschungsgemeinschaft (DFG), and by J.G. from the VW Stiftung. “Big Data in den Lebenswissenschaften der Zukunft”, Nr. A129197.

## Author contributions

S.B. designed the project. S.B. and K.R. made the figures and wrote the manuscript.

A.H. and D.M. performed the experiments. G.S. and J.G. managed and coordinated RNA seq analysis. All authors commented on the manuscript.

---

**Supplementary Figure 1.**
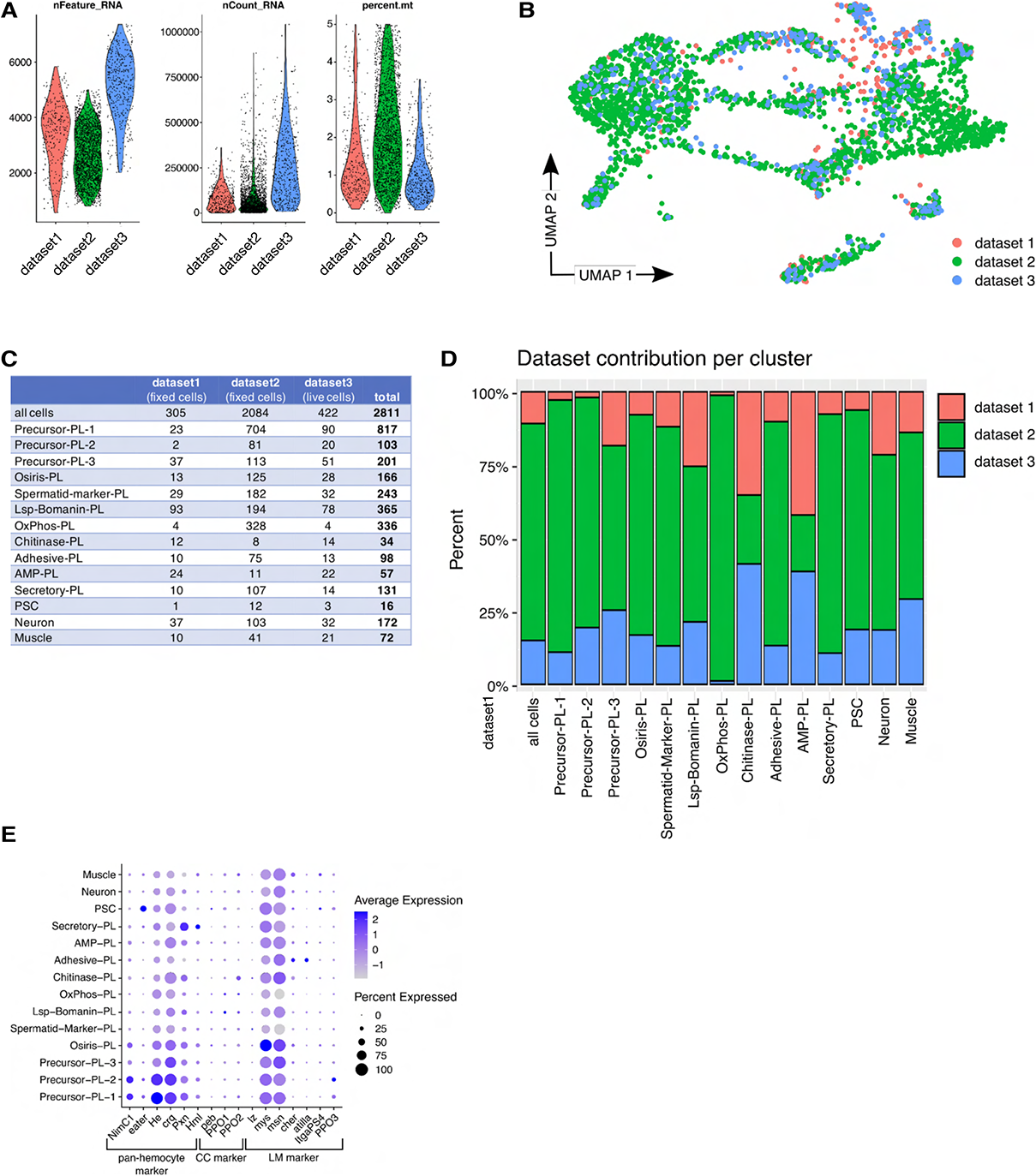
Violin plots representing the number of Features, UMI counts and percent of mitochondrial genes per cell and per dataset. B) UMAP plot with cells labeled by dataset origin. c, d) Contribution of cells per dataset in a table (C) or represented as a barplot (D). Note that all datasets (two fixed and one live cell dataset) contribute to every cluster. E) Average expression and percent expression per cluster of known pan-hemocyte, crystal cell and lamellocyte markers.

**Supplementary Figure 2.**
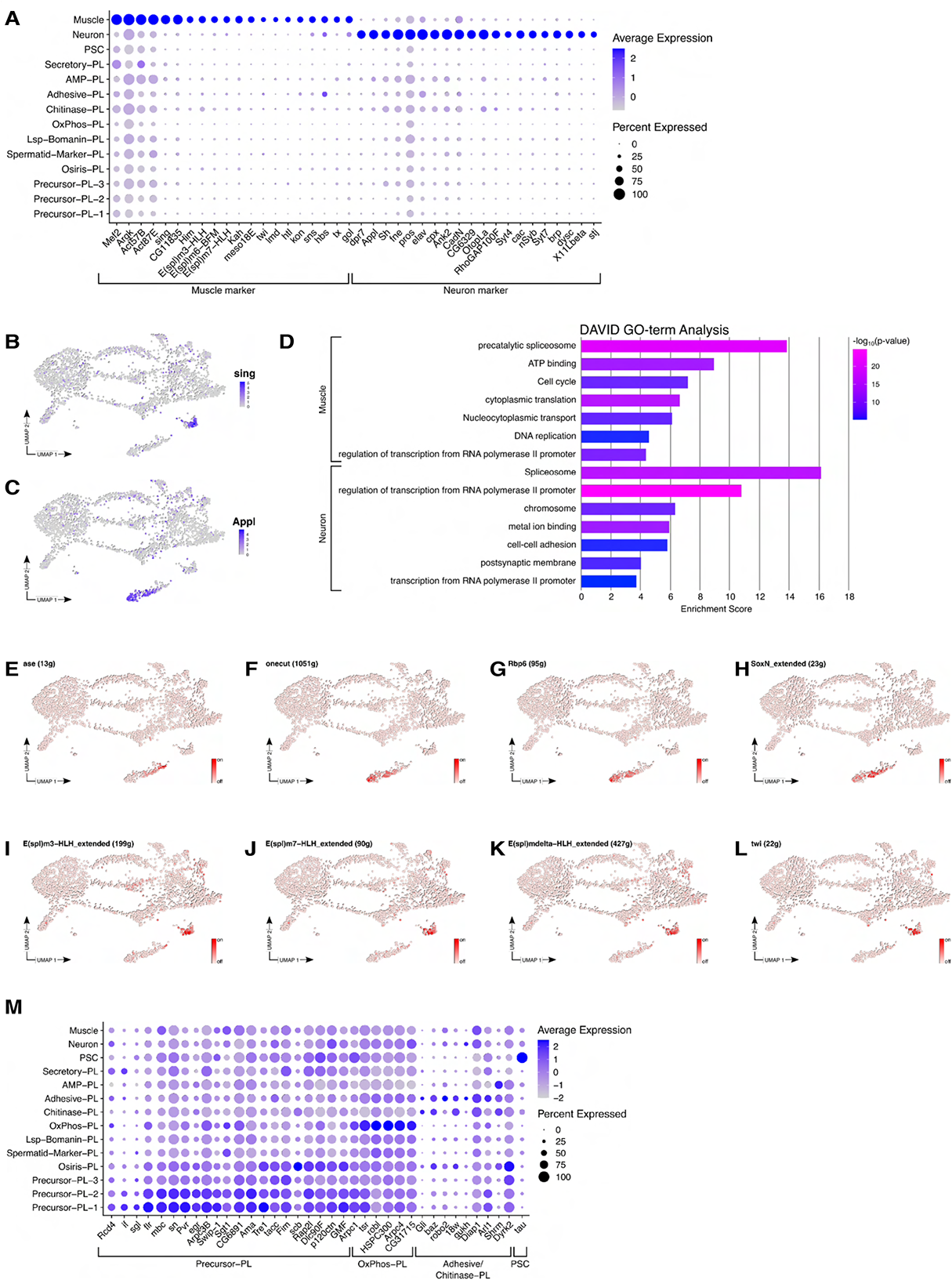
Dotplot showing the average expression and percent expression of muscle and neuron markers. B-C) Expression of the muscle marker *sing* (B) and the neuron marker *Appl* (C) on the UMAP plot. D) DAVID 2021 GO-term analysis of Neuron and Muscle markers. For each of the top seven annotation cluster a representative term is presented. Barplot shows enrichment score of the annotation cluster and p-value of the GO-term. E-L) SCENIC determined activity of transcription factors active in Neuron (E-H) or Muscle (I-L) on the UMAP plot. E) *ase*, F) *onecut*, G) *Rbp6*, H) *SoxN*, I) *E^20^m3-HLH*, J) *E^20^m7-HLH*, K) *E^20^mdelta-HLH*, L) *twi*. M) Average expression and percent expression per cluster of cytoskeleton and cell motility genes identified in a bulk RNA-sequencing experiment comparing larval and pupal hemocytes.

**Supplementary Figure 3.**
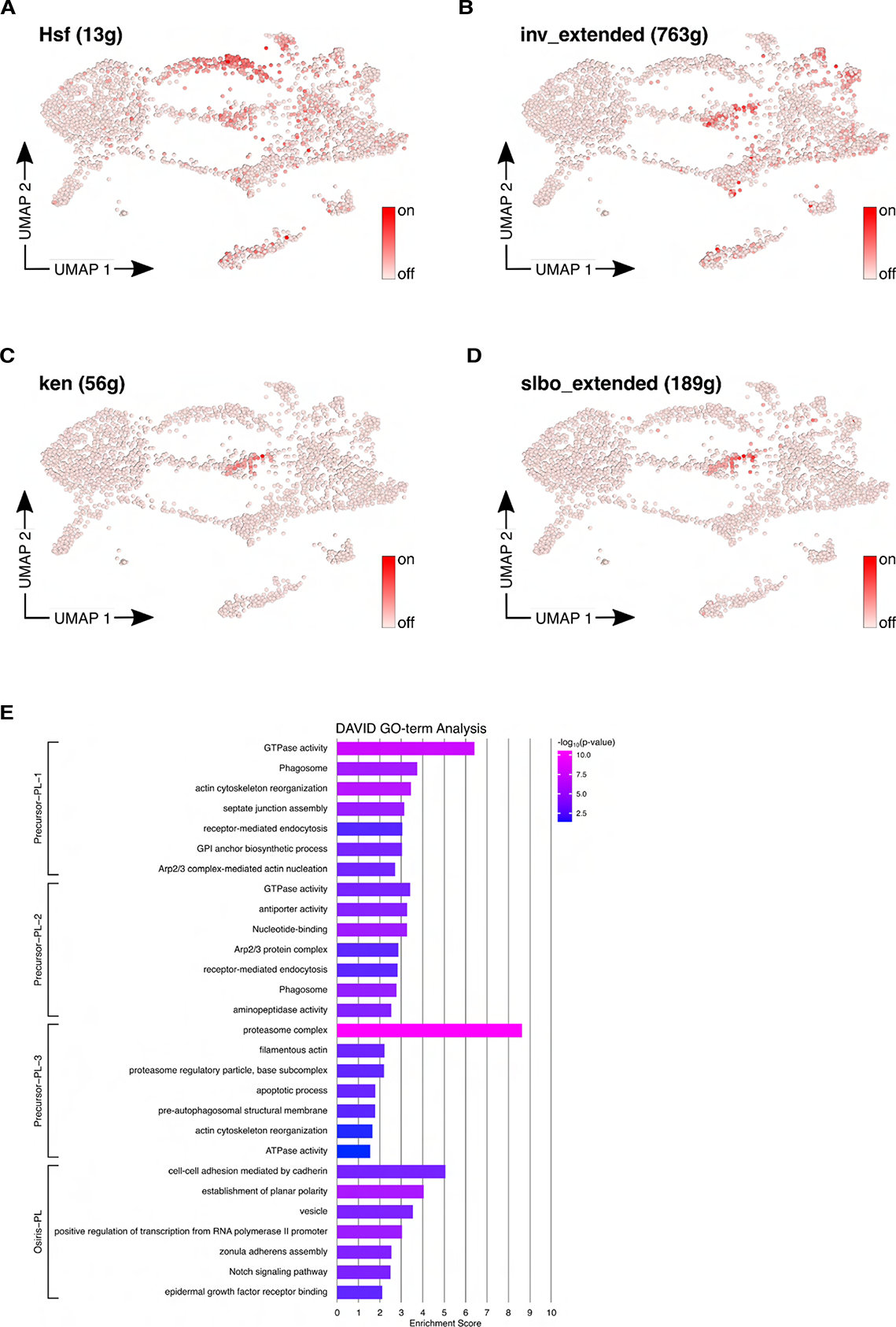
A-D) Activity of transcription factors on the UMAP plot as identified by SCENIC. A) *Hsf*, A) *inv*, C) *ken*, D) *slbo*. E) Marker genes of Precursor-PL-1 – 3 or Osiris-PL were analysed with DAVID 2021. For the top seven annotation cluster one representative GO-term is shown. Barplot shows enrichment score of the annotation cluster and is colored by GO-term p-value.

**Supplementary Figure 4.**
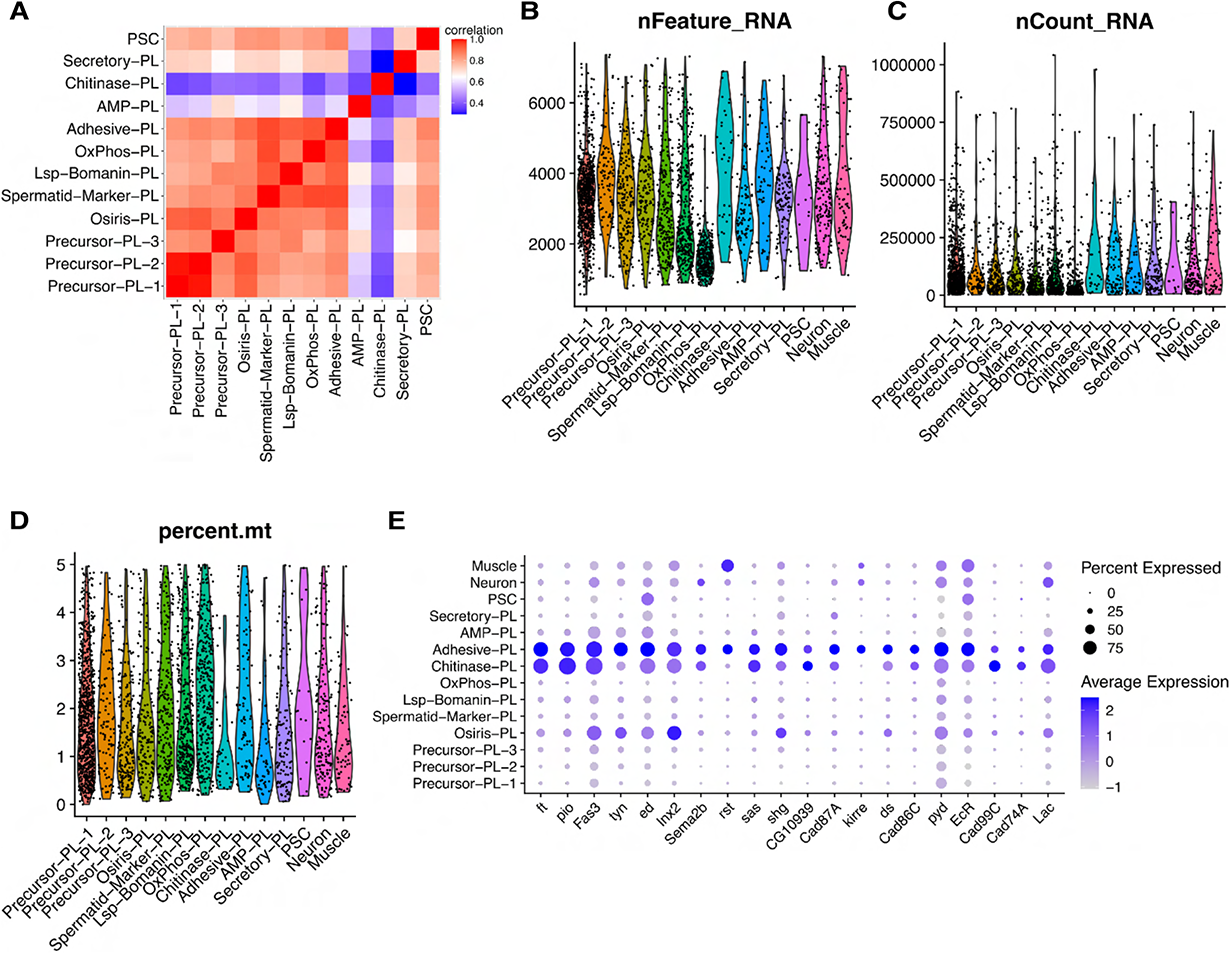
A) Cross cluster correlation analysis of hemocyte cell types. *srp*^high^ clusters correlate well with each other and with several differentiated plasmatocyte types. Transcriptomes of plasmatocytes Chitinase-PL and AMP-PL differ dramatically, suggesting a high level of specification. b-d) Violin plot showing the number of (B) features, (C) UMIs or (D) percent of mitochondrial genes per cluster. E) Average and percent expression of genes implicated in cell adhesion identified as markers for Adhesive-PL. Note that most genes are also expressed in Chitinase-PL.

**Supplementary Figure 5.**
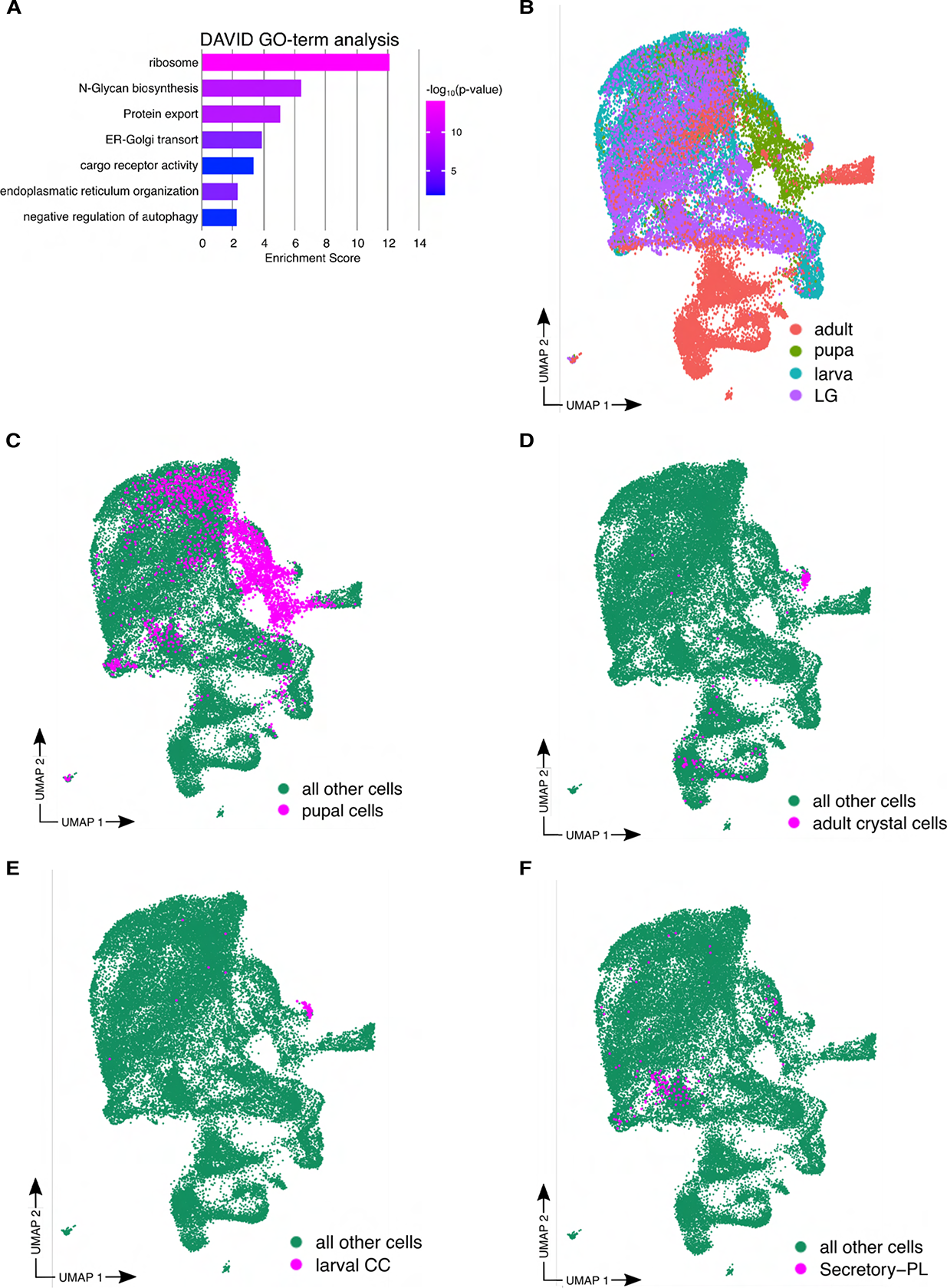
A) GO-term analysis of Secretory-PL markers performed with DAVID 2021. Representative GO-terms for the top seven annotation clusters are shown. Enrichment score of annotation clusters and p-value of GO-terms are presented in A barplot. B-F) UMAP plots of integrated hemocyte datasets across different developmental stages. B) Labelled by dataset. C) Pupal cells are highlighted in magenta. D) Adult crystal cells are highlighted in magenta. E) Larval crystal cells are shown in magenta. F) Pupal Secretory-PL are represented in magenta.

**Supplementary Figure 6.**
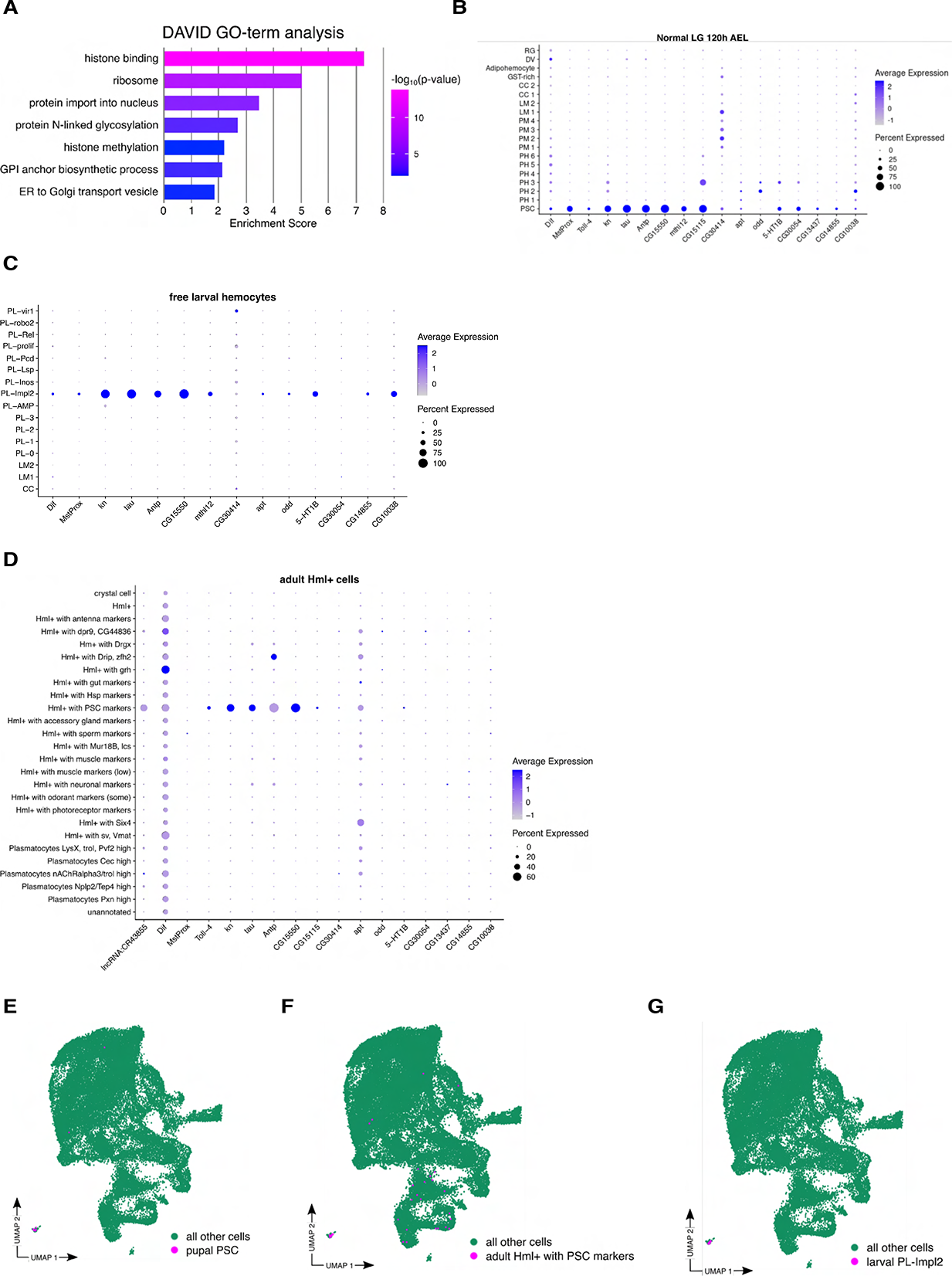
A) PSC markers were analysed with DAVID 2021 and representative GO-terms of the top seven GO-terms are presented. Barplot shows the enrichment score of the annotation cluster and is colored by p-value of the GO-term. B-D) Dotplots of markers of pupal PSCs showing average expression and percent expression per cluster in B) the normal lymph gland ^20^ at 120h afer egg laying, C) free larval hemocytes of embryonic origin and D) in adult Hml positive cells. D-F) UMAP plot of integrated datasets across developmental stages showing the location of D) pupal PSC, E) adult Hml positive cells with PSC markers and F) larval PL-Impl2 cells.

## Supplementary movies

**Supplementary movie 1. PSC cells randomly migrate in prepupae**.

Representative spinning disc microscopy video of randomly migrating pupal *hml+* (left) versus *kn+* (right) cells expressing an EGFP transgene imaged from a living prepupa (4h APF). Note the difference of *hml+* hemocytes marked by a yellow arrow. Scale bar represents 20 µm. Cells were imaged for 30 minutes.

**Supplementary movie 2. PSC cells are wound responsive and differ in size compared to *hml-*marked plasmatocytes**

Representative spinning disc microscopy videos of *hml+* (left) and *kn+* (right) cells in the Drosophila abdomen 16h APF upon single cell laser ablation. Cells are imaged for 60 minutes after ablation in a 30 seconds interval and tracked afterwards using Imaris. Representative migratory tracks are shown (colored, jagged lines). Characteristic plasmatocytes are marked by yellow arrows compared to a small, spiky PSC cell marked by a white arrow. Yellow asterisk marks the ablated wounding site. Scale bar represents 20 µm.

**Supplementary movie 3. PSC cells switch from random to directed migration upon single cell laser ablation.**

Representative spinning disc microscopy video with a magnification (indicated by yellow rectangle) of single *kn+* cells that migrate towards a laser-ablated cell marked by a yellow asterisk. Note the difference between plasmatocytes marked by yellow arrows and PSC cells. Scale bar represent 20 µm.

**Supplementary Data 1**

List of differentially expressed genes across all 14 clusters identified with Seurat v4.1.1 FindAllMarkers() command.

**Supplementary Data 2**

Data associated with Figure 1C. This list contains the top50 genes identified with Seurat v4.1.1 FindAllMarkers() and is an abbreviated version of Supplementary Data 1.

**Supplementary Data 3**

Differentially expressed genes of Precursor-PL-1 identified with Seurat v.1.1 FindMarkers() used for DAVID 2021 GO-term analysis.

**Supplementary Data 4**

Differentially expressed genes of Precursor-PL-2 identified with Seurat v.1.1 FindMarkers() used for DAVID 2021 GO-term analysis.

**Supplementary Data 5**

Differentially expressed genes of Precursor-PL-3 identified with Seurat v.1.1 FindMarkers() used for DAVID 2021 GO-term analysis.

**Supplementary Data 6**

Differentially expressed genes of Osiris-PL identified with Seurat v.1.1 FindMarkers() used for DAVID 2021 GO-term analysis.

**Supplementary Data 7**

Differentially expressed genes of Spermatid-Marker-PL identified with Seurat v.1.1 FindMarkers() used for DAVID 2021 GO-term analysis.

**Supplementary Data 8**

Differentially expressed genes of Lsp-Bomanin-PL identified with Seurat v.1.1 FindMarkers() used for DAVID 2021 GO-term analysis.

**Supplementary Data 9**

Differentially expressed genes of OxPhos-PL identified with Seurat v.1.1 FindMarkers() used for DAVID 2021 GO-term analysis.

**Supplementary Data 10**

Differentially expressed genes of Chitinase-PL identified with Seurat v.1.1 FindMarkers() used for DAVID 2021 GO-term analysis.

**Supplementary Data 11**

Differentially expressed genes of Adhesive-PL identified with Seurat v.1.1 FindMarkers() used for DAVID 2021 GO-term analysis.

**Supplementary Data 12**

Differentially expressed genes of AMP-PL identified with Seurat v.1.1 FindMarkers() used for DAVID 2021 GO-term analysis.

**Supplementary Data 13**

Differentially expressed genes of Secretory-PL identified with Seurat v.1.1 FindMarkers() used for DAVID 2021 GO-term analysis.

**Supplementary Data 14**

Differentially expressed genes of PSC identified with Seurat v.1.1 FindMarkers() used for DAVID 2021 GO-term analysis.

**Supplementary Data 15**

Differentially expressed genes of Neuron identified with Seurat v.1.1 FindMarkers() used for DAVID 2021 GO-term analysis.

**Supplementary Data 16**

Differentially expressed genes of Muscle identified with Seurat v.1.1 FindMarkers() used for DAVID 2021 GO-term analysis.

**Supplementary Data 17**

Scaled regulon activity identified by SCENIC v1.3.1. A subset of this data is visually presented in Figure 1D.

**Supplementary Data 18**

Top active transcription factors per cell type identified by SCENIC v1.3.1.

**Supplementary Data 19**

Validation of marker genes for each cluster of *Drosophila* hemocytes. A list of Gal4-enhancer trap and GFP-exon trap fly lines used for ex vivo validation is included.

**Script 1**

R script for data processing steps including quality filtering, normalization, data integration, clustering and cluster annotation using the Seurat package.

**Script 2**

R script for the pseudotime analysis using Monocle3

**Script 3**

R script for the identification of transcription factor activity with SCENIC.

**Script 4**

R script for the integration of datasets from different developmental stages using the Seurat batch correction algorithm.

**Script 5**

R script containing an overview of how the processed data can be read into R and analyzed for gene expression.

